# Brain imaging correlates of general intelligence in UK Biobank

**DOI:** 10.1101/599472

**Authors:** SR Cox, SJ Ritchie, C Fawns-Ritchie, EM Tucker-Drob, IJ Deary

## Abstract

The associations between indices of brain structure and measured intelligence are not clear. In part, this is because the evidence to date comes from mostly small and heterogenous studies. Here, we report brain structure-intelligence associations on a large sample from the UK Biobank study. The overall N = 29,004, with N = 18,363 participants providing both brain MRI and cognitive data, and a minimum N = 7318 providing the MRI data alongside a complete four-test battery. Participants’ age range was 44-81 years (M = 63.13, SD = 7.48). A general factor of intelligence (*g*) was extracted from four varied cognitive tests, accounting for one third of the variance in the cognitive test scores. The association between (age-and sex-corrected) total brain volume and a latent factor of general intelligence is *r* = 0.275, 95% C.I. = [0.252, 0.299]. A model that incorporated multiple global measures of grey and white matter macro-and microstructure accounted for more than double the *g* variance in older participants compared to those in middle-age (13.4% and 5.9%, respectively). There were no sex differences in the magnitude of associations between *g* and total brain volume or other global aspects of brain structure. The largest brain regional correlates of *g* were volumes of the insula, frontal, anterior/superior and medial temporal, posterior and paracingulate, lateral occipital cortices, thalamic volume, and the white matter microstructure of thalamic and association fibres, and of the forceps minor.

## 1. Introduction

The association between brain volume and intelligence has been one of the most regularly-studied—though still controversial—questions in cognitive neuroscience research. The conclusion of multiple previous meta-analyses is that the relation between these two quantities is positive and highly replicable, though modest (McDaniel, 2005; Gignac & Bates, 2017; Pietschnig, Penke, Wicherts, Zeiler, & Voracek, 2015), yet its magnitude remains the subject of debate. The most recent meta-analysis, which included a total sample size of 8,036 participants with measures of both brain volume and intelligence, estimated the correlation at *r* = 0.24 (Pietschnig et al., 2015). A more recent re-analysis of the meta-analytic data, only including healthy adult samples (N = 1,758), found a correlation of *r* = 0.31 (Gignac & Bates, 2017). Furthermore, the correlation increased as a function of intelligence measurement quality: studies with better-quality intelligence tests—for instance, those including multiple measures and a longer testing time—tended to produce even higher correlations with brain volume (up to 0.39). In a meta-analysis, issues of cross-cohort heterogeneity might have an important bearing on the magnitude of the correlation.

Here, we report an analysis of data from a large, single sample with high-quality MRI measurements and four diverse cognitive tests. We use latent variable modelling to create a general intelligence (‘*g*’) factor from the cognitive test and estimate its association with both total brain volume and several more fine-grain imaging-derived indices of brain structure. We judge that the large N, study homogeneity, and diversity of cognitive tests relative to previous large scale analyses provides important new evidence on the size of the brain structure-intelligence correlation. By investigating the relations between general intelligence and characteristics of many specific regions and subregions of the brain in this large single sample, we substantially exceed the scope of previous meta-analytic work in this area.

There is considerable debate about what the association between brain size and general intelligence means. It is unclear, for example, whether brain size is a direct proxy for neuron number (discussed in Pietschnig et al., 2015). There is also an apparent paradox that there are substantial sex differences in total brain volume (on the order of 1.41 standard deviations; Ritchie et al., 2018) but no sex differences in mean intelligence (Johnson et al., 2009). More recent work indicates that multiple brain properties might be required to better explain individual differences in general intelligence, and some of these might be compensatory for differences in overall brain size (Luders et al., 2004; Deary et al., 2010; Kievit et al., 2012, 2014; Ritchie et al., 2015). For example, in an older cohort with a narrow age range (N = 672), Ritchie et al. (2015) found that incorporating multiple global, but tissue-specific, brain MRI measures (including tissue volumes, measures of white matter microstructure, and hallmarks of brain ageing) accounted for up to 21% of the variance in general intelligence, which was substantially higher than could be accounted for by total brain size alone (∼12%).

Such results, combined with indications that there is regional heterogeneity in the magnitude of intelligence associations across both grey and white matter (Jung & Haier., 2007; Deary et al., 2010; Basten et al., 2015; Karama et al., 2011; Cox et al., 2018; Ryman et al., 2016), extend the focus beyond a single, well-replicated proxy (total brain volume) and toward tissue-(and region-) specific associations with general intelligence. One of the most influential accounts of the neurobiological underpinnings of general intelligence (also known as general intelligence or “*g*”) has been the Parieto-Frontal Integration Theory (P-FIT; Jung & Haier, 2007). The P-FIT was initially based on a synthesis of disparate structural and functional brain imaging results. However, none of the Brodmann regions implicated in intelligence were supported by more than 60% of the studies reviewed, which the authors pointed out might be considered a relatively weak consensus.

The P-FIT model implicates, the following regions as being associated with intelligence differences: the lateral frontal, superior temporal, medial temporal, parietal and extrastriate (lateral occipital) regions, along with the white matter tracts that connect them. Specific reference was originally made to the arcuate fasciculus; this pathway is variously described as being just adjacent to the superior longitudinal fasciculus, or as one of the components thereof (Kamali et al., 2014; Dick and Tremblay, 2012). Together, these form a ‘dorsal stream’ of anterior-posterior cortical connectivity. Alongside other fibres such as the inferior longitudinal, inferior fronto-occipital, uncinate, and cingulum fasciculi, these ‘association’ fibres—along with the genu of the corpus callosum (forceps minor)—facilitate connectivity across the distal cortical regions highlighted by the P-FIT model. The model has generally received support from subsequent work (Deary et al., 2010; Basten et al., 2015; Karama et al., 2011; Cox et al., 2018; Ryman et al., 2016).

As with the meta-analyses on brain volume and intelligence described above (Pietschnig et al., 2015, Gignac and Bates, 2017), the broad heterogeneity of studies on the P-FIT might produce a less precise picture of the brain basis of cognitive abilities. Further evaluation of the model would greatly benefit from large-sample research that investigates the grey and white matter components of this putative intelligence framework, together in the same analysis. We conduct that analysis in the present study.

We capitalize on data from the UK Biobank study, a large-scale biomedical study of health and wellbeing, which includes brain MRI and various measures of cognitive ability. The UK Biobank participants have completed various cognitive measures; originally, they were administered a battery of bespoke tests with relatively poor reliability (Lyall et al., 2016). Using an earlier data release (Ritchie et al., 2018), we previously estimated the correlation between brain size and one of those tests, “Fluid Intelligence” (which we refer to as Verbal-Numerical Reasoning) to be *r* = 0.177. We found that the correlation did not differ by sex. Another study using an earlier release of UK Biobank imaging data examined the association between Verbal-Numerical Reasoning and brain size, reporting a correlation of *r* = 0.19 (N = 13,608; Nave et al., 2018). In addition, analyses of regional white and 10 grey matter measures have been reported with respect to Verbal-Numerical Reasoning in an earlier UK Biobank release; however, the authors of that study cited several reasons to doubt that this test, in isolation, is a valid indicator of fluid cognitive ability (Kievit et al., 2018; see also Hagenaars et al., 2016).

In this pre-registered study, we use a newer subset of UK Biobank participants who have completed an enhanced cognitive assessment battery at their brain imaging assessment. The overlap of the complete cognitive battery and the various MRI measures ranges from N = 8165 to 7318 following exclusions that are described below. Their data have only recently been released, and have not previously been analysed by our team. The enhanced cognitive battery includes three new measures based on standardised cognitive tests: Symbol-Digit Substitution, Matrix Reasoning, and Trail-Making. These three tests, combined with the previous Verbal-Numerical Reasoning measure, allows the estimation of brain imaging associations with a latent factor of general intelligence (*g*), that arguably gives coverage of the cognitive domains of reasoning, processing speed, working memory, and executive function. In a large sample size, the current study design thus: results in a better-quality cognitive measure than was previously possible in the UK Biobank data; mitigates variability in the administration and measurement of cognitive and brain imaging constructs (potentially allowing for stronger brain-intelligence correlations; Gignac & Bates, 2017); and, given the detailed brain imaging measures available, facilitates a detailed estimate of the global and regional brain correlates of latent general intelligence.

Our analyses followed a preregistered protocol and 4 hypotheses (https://osf.io/w7evd/). First, we tested whether the four cognitive tests were correlated moderately-highly (*r* > 0.40), and formed a latent general factor that explains 40% or higher of the variance across tests. Next, we examined the association between this cognitive factor and total brain volume, and we hypothesised that there would be no significant sex difference in the size of the brain-cognitive correlation for any of the models. We then hypothesised that different global measures of grey and white matter would each account for significant unique variance in *g*. We then aimed to test associations between general intelligence and brain grey and white matter regional measures, hypothesising that the strongest associations would concur with regions implicated by the Parieto-Frontal Integration Theory of intelligence (P-FIT; Jung & Haier, 2007).

## 2. Methods

The UK Biobank study is a large-scale biomedical study of health and wellbeing, which includes brain MRI and measures of cognitive function (Sudlow et al., 2015). Cognitive tests and brain imaging data were acquired on the same assessment day. The tests used here were administered at the UK Biobank brain imaging assessment. The imaging assessment took place at 3 different assessment centres. The majority were in Manchester, with more recent appointments now also taking place in Newcastle, and most recently in Reading. Cognitive tests were administered to participants working independently on a touchscreen computer with no tester observing. MRI data was acquired at the three sites using the same hardware and software. The current data release from UK Biobank initially included 30,316 participants who attended the scanning appointment, i.e. they had a record for age at scanning. Following exclusions, the total N = 29,004. The minimal N with complete cognitive-MRI overlap was N = 7318; further information is provided in Statistical Analysis, Table 1 and Figures 1 and S1. UK Biobank Field IDs are listed in Table S1. Most of them had scores for the Verbal-Numerical Reasoning test (always administered at the MRI appointment). Many fewer had the more recently-introduced cognitive tests (Matrix Pattern Completion, Symbol-Digit and Trail Making Test Part B). Missing MRI data was due to the lag between MRI acquisition and its subsequent processing for release by the UK Biobank. All data and materials are available via UK Biobank (http://www.ukbiobank.ac.uk).

**Table 1.** Participant characteristics split out by assessment centre, and for the full sample.

|  | Manchester |  | Reading |  | Newcastle |  | Full Sample |  |
| --- | --- | --- | --- | --- | --- | --- | --- | --- |
|  | M(SD)<sup>a</sup> | N | M(SD) | N | M(SD) | N | M(SD) | N |
| **Age (years)** | 62.818 (7.484) | 22037 | 65.202 (7.299) | 866 | 63.962 (7.379) | 6101 | 63.130 (7.480) | 29004 |
| **Sex (F:M)** | 11315 : 10722 | 22037 | 453 : 413 | 866 | 3256 : 2845 | 6101 | 15024 : 13980 | 29004 |
| **Matrix Reasoning** | 8.047 (2.113) | 8994 | 8.277 (2.045) | 850 | 7.904 (2.124) | 5671 | 8.007 (2.115) | 15515 |
| **Symbol-Digit** | 19.212 (5.255) | 9019 | 18.753 (5.152) | 858 | 18.714 (5.235) | 5681 | 19.005 (5.247) | 15558 |
| **VNR** | 6.778 (2.070) | 20455 | 6.800 (2.058) | 860 | 6.466 (2.001) | 5668 | 6.713 (2.061) | 26983 |
| **TMTb<sup>a</sup>** | 496 (236) | 9103 | 512 (230) | 866 | 503(236) | 5710 | 499 (235) | 15679 |
| **TBV** | 1170062 (110900) | 17223 | - | 0 | 1154425 (110376) | 3104 | 1167674 (110960) | 20327 |
| **GM** | 618392 (55474) | 17226 | - | 0 | 611240 (54923) | 3104 | 617300 (55448) | 20330 |
| **NAWM** | 547110 (61736) | 16146 | - | 0 | 537269 (61090) | 3062 | 545541 (61737) | 19208 |
| **WMH<sup>a</sup>** | 2500 (3641) | 16146 | - | 0 | 3290(4506) | 3062 | 2622 (3816) | 19208 |
| **gFA** | 0.025 (0.824) | 15448 | - | 0 | -0.131 (0.834) | 2989 | 0.00 (0.827) | 18437 |
| **gMD** | 0.022 (0.918) | 15448 | - | 0 | -0.113 (0.907) | 2989 | 0.00 (0.918) | 18437 |

*Note.* Means and standard deviations (SD) reported, except for <sup>a</sup>median and interquartile ranges are given. VNR: verbal numerical reasoning, TMTb: Trail Making Test Part b, TBV: total brain volume, GM: grey matter volume, NAWM: normal-appearing white matter volume, WMH: white matter hyperintensity volume, gFA: general factor of white matter fractional anisotropy, gMD: general factor of white matter mean diffusivity.

**Figure 1.**
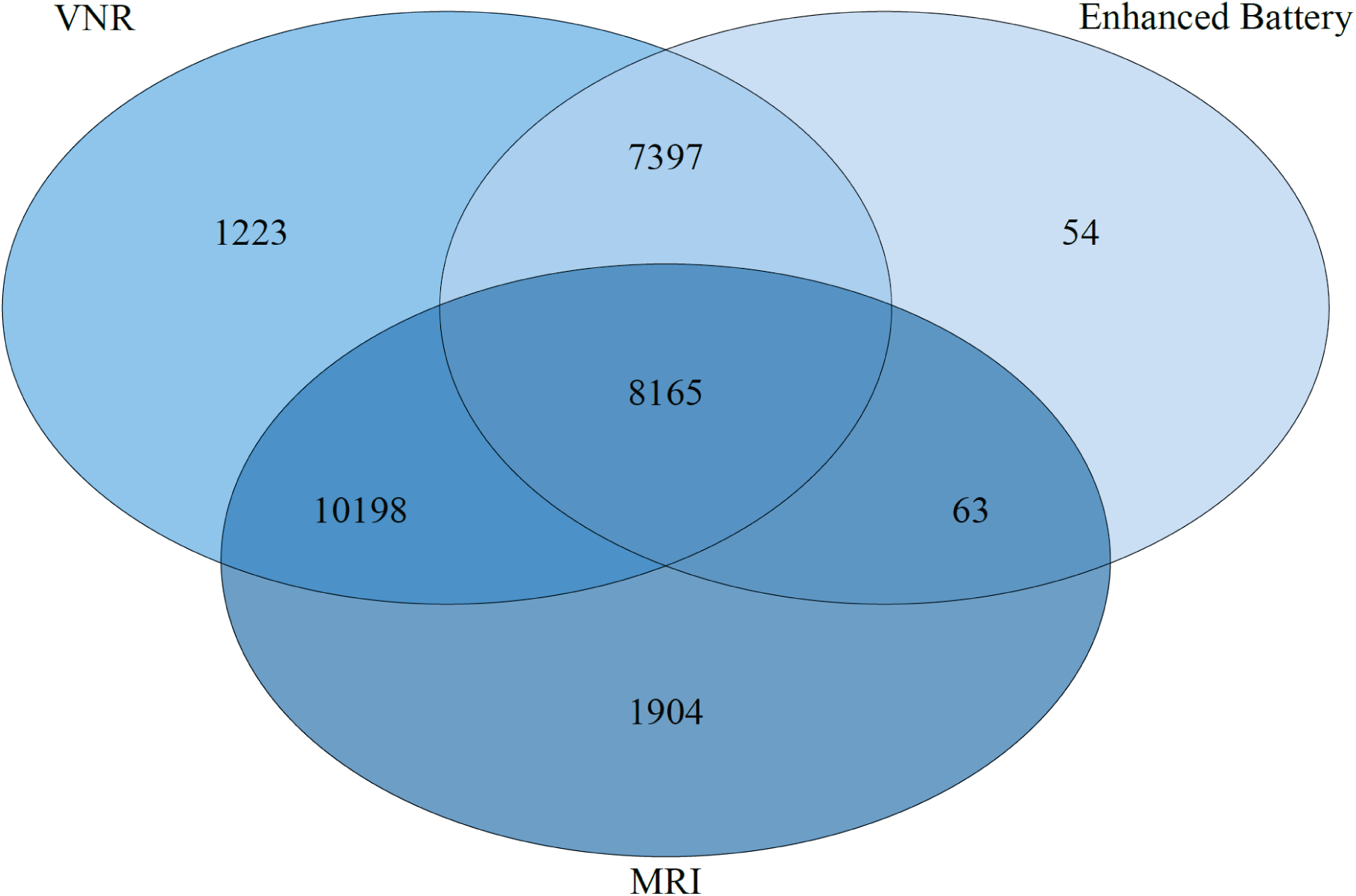
Overlap between initial cognitive measures in the imaging visit (VNR; verbal numerical reasoning), MRI measures, and the Enhanced Cognitive Battery (Matrix Reasoning, Symbol-Digit and Trail-Making Part B) among the 29,004 participants included in the current analysis. For ease of illustration, the MRI numbers are based on grey matter volume (highest N among global MRI measures), and the Enhanced Battery numbers are based upon Trail Making Part B (highest N among the Enhanced Battery). A total of 18,363 have MRI and at least one cognitive test. There are slight variations in missingness among Enhanced Battery and MRI measures (see Table 1); such that the precise overlap among *all* cognitive tests and grey matter volume is 8124, and this becomes lower as a function of the availability of brain imaging measures (minimal overlap = 7318 with dMRI data).

### 2.1 Cognitive Tests

The four cognitive tests used in the current study were: Matrix Reasoning, Symbol-Digit Substitution, Verbal-Numerical Reasoning, and Trail-Making Test. Specifically, for the Trail-Making Test, we used part B, since this test includes both elements of speed and executive functioning (Salthouse, 2011).

#### Matrix Pattern Completion

The UK Biobank Matrix Reasoning test is an adapted version of the Matrices test in the COGNITO battery (Ritchie et al., 2014). This test of non-verbal fluid reasoning requires participants to inspect a grid pattern with a piece missing in the lower right-hand corner and select which of the multiple choice options at the bottom of the screen completes the pattern both horizontally and vertically. This 15-item test assesses participants’ ability to problem solve using novel and abstract materials. The score is the number of correctly answered questions in three minutes.

#### Symbol-Digit Substitution

was used as a measure of processing speed. It is similar in format to the Symbol Digit Modalities Test (Smith, 1991), which is a well-validated measure of processing speed. At the top of the screen, participants were shown a key pairing shapes with numbers. Beneath the key were rows of shapes with an empty box under each shape. Using the key, participants had 60 seconds to enter the number in the empty boxes that are paired with the shapes. Participants were instructed to work as quickly and as accurately as possible. The score is the number of correct symbol-digit matches made in 60 seconds.

#### *Verbal-Numerical Reasoning* (referred to as “Fluid Intelligence” in UK Biobank)

Participants were presented with 13 multiple choice questions assessing verbal and numerical reasoning abilities. The score is the number of questions answered correctly in 2 minutes.

#### Trail-Making Test Part B

This test is a computerised version of the Halstead-Reitan Trail-Making Test (Reitan & Wolfson, 1985). It is often said to be an assessment of executive function. In part B, participants were presented with the numbers 1-13, and the letters A-L arranged quasi-randomly on a computer screen. The participants were instructed to switch between touching the numbers in sequential order, and the letters in alphabetical order (e.g., 1-A-2-B-3-C) as quickly as possible. The score is the time (in deci-seconds) taken to successfully complete the test. Those with a score coded as 0 (denoting “Trail not completed”) had their score set to missing.

### 2.2 Brain Imaging Acquisition and Analysis

All brain MRI data were acquired on a Siemens Skyra 3T scanner with a standard Siemens 32-channel head coil, in accordance with the open-access protocol (http://www.fmrib.ox.ac.uk/ukbiobank/protocol/V4_23092014.pdf), documentation (http://biobank.ctsu.ox.ac.uk/crystal/docs/brain_mri.pdf), and publication (Alfaro-Almagro et al., 2018). T_1_-weighted MPRAGE data was acquired in the sagittal plane at 1mm isotropic resolution; the T_2_-weighted FLAIR acquisition at 1.05 × 1 × 1 mm resolution, was also acquired in the sagittal plane. The diffusion MRI (dMRI) data was acquired using a spin-echo echo-planar sequence with 10 T_2_-weighted (b ≈ 0 s mm^2^) baseline volumes, 50 b = 1000 s mm^-2^ and 50 b = 2000 s mm^-2^ diffusion-weighted volumes, with 100 distinct diffusion-encoding directions and 2 mm isotropic voxels. We used global and regional brain Imaging Derived Phenotypes (IDPs) provided by the UK Biobank brain imaging team: total brain volume (TBV, which is the sum of grey and white matter and excludes cerebrospinal fluid), grey matter volume (GM), and white matter volume (WM) from FSL FAST (Zhang et al., 2001), 14 subcortical volumes using FSL FIRST (Patenaude et al., 2011) and white matter hyperintensity volume (WMH) using BIANCA (Griffanti et al., 2016), which uses both T1-weighted and T2-weighted volumes. We estimated normal-appearing white matter volume (NAWMV) as the difference between total WMV and WMHV. Regional brain information was also available as UK Biobank IDPs in the form of tract-averaged fractional anisotropy and mean diffusivity for each of 27 white matter tracts using AutoPtx (de Groot et al., 2013), and as individual grey matter cortical segmentations, derived using FSL FAST, using the Harvard-Oxford Atlas (https://fsl.fmrib.ox.ac.uk/fsl/fslwiki/Atlases). These white matter tracts and cortical regions are shown in Figure 2.

**Figure 2.**
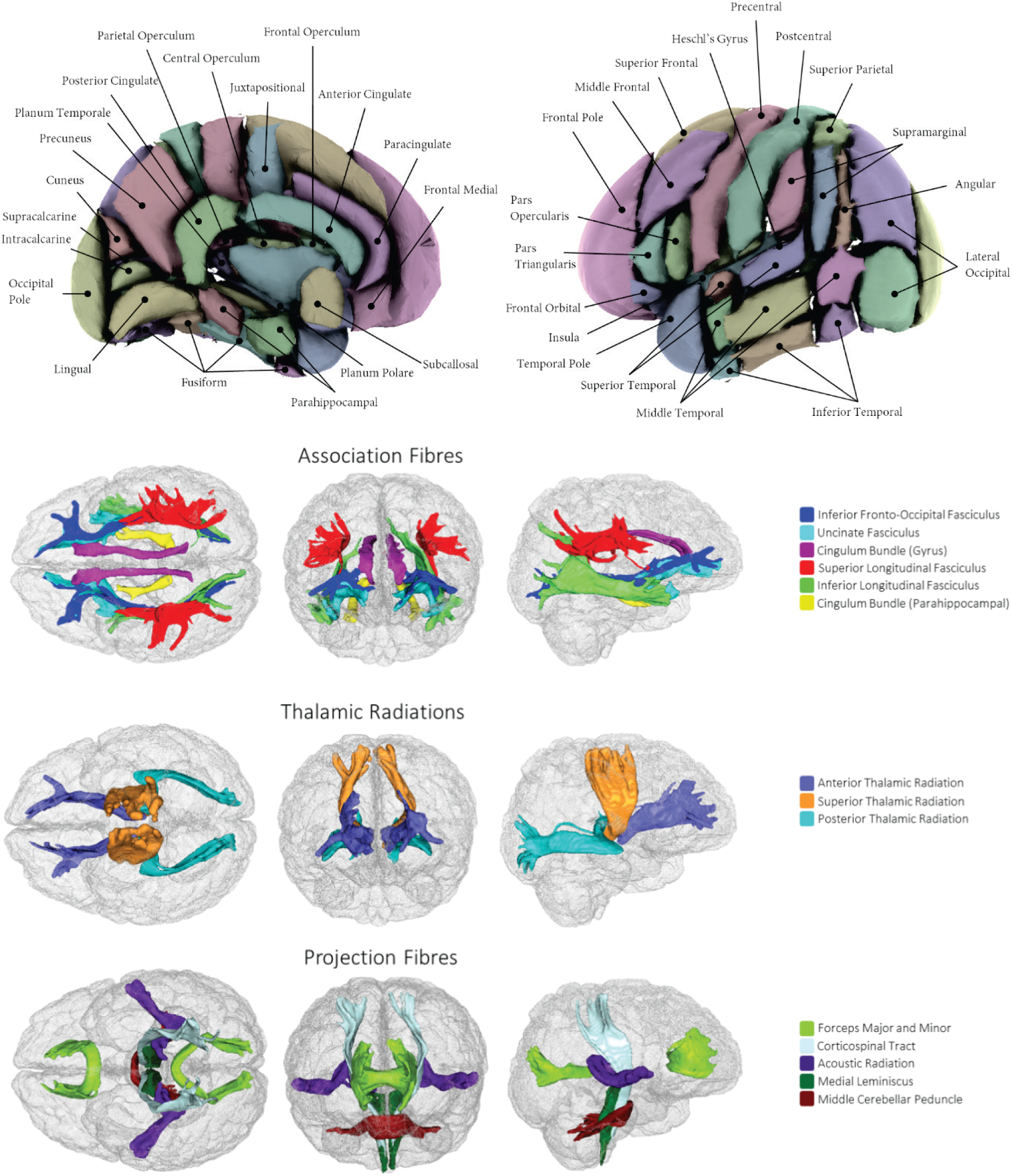
Brain imaging regions of interest according to the Harvard-Oxford Atlas (**top: cortical regions**) and AutoPtx (**bottom: white matter tracts**; adapated from Cox et al., 2019).

### 2.3 Statistical Analyses

Outliers (+/-4SDs) were removed from brain and cognitive measures. All individuals who reported any of the neurological or neurodegenerative health conditions listed in Supplementary Material (Table S1) were removed prior to analysis. All participants were of White European ancestry. Of an initial 30,316 of the UK Biobank participants who had a record of age at attending the imaging assessment, 29,004 provided data on at least one of the primary variables of interest (global brain imaging or cognitive) following exclusions, and 18,363 had at least one cognitive test and MRI data. All structural equation modelling (SEM) analyses were conducted using Full Information Maximum Likelihood (FIML) estimation within R, using the *lavaan* package for SEM (Rosseel, 2012). Throughout, indicators were corrected for age and sex, with the MRI variables also corrected for scanner head positioning confounds (X, Y and Z coordinates provided by the UK Biobank team: UKB IDs: 25756, 25757, 25758). In contrast to our pre-registration, covariates (age, sex, MRI confounds) were applied to manifest variables within SEMs (i.e. we did not need to residualise data outside the model to enable model convergence and fit). FIML takes advantage of all available data, including data from participants who are missing data on some of the dependent variables. Model fits were assessed with a chi-squared test, Root Mean Square Error of Approximation (RMSEA), the Comparative Fit Index (CFI), the Tucker-Lewis Index (TLI), and the Standardized Root Mean Square Residual (SRMR).

#### 2.3.1 Estimating a latent general factor of general intelligence, ‘g’

We performed a confirmatory factor analysis (CFA) of the 4 cognitive tests: Symbol-Digit Substitution, Matrix Reasoning, Trail-Making Test Part B, and Verbal-Numerical Reasoning. We hypothesised that the four tests would correlate moderately-highly (with intercorrelations of r > .40), and would form a single latent general factor explaining ∼40% of the variance across the 4 tests, with good fit to the data (CFI and TLI > 0.95, SRMR and RMSEA <0.05). We ran a version without, and then with age and sex correction at the manifest level. Since principal components analyses (PCAs) are commonly also used in intelligence research (e.g. Nave et al., 2018), but do not separate common and test-specific variance, we also provide a PCA estimate of *g* (tests not corrected for age and sex) using the first unrotated principal component, for comparison with the CFA.

#### 2.3.2 Associations between general intelligence (g) and global brain MRI measures

Next, we estimated the association of the latent general intelligence factor (‘*g*’) with total brain volume, and then 6 global measures of grey and white matter: grey matter, normal-appearing white matter and white matter hyperintensity volumes (TBV, GM, NAWM, WMH), and general factors of fractional anisotropy and mean diffusivity (*g*FA and *g*MD). The general factors of white matter microstructure were formed from diffusion indices for white matter pathways of interest, which were extracted from confirmatory factor analysis, as previously described (Cox et al., 2016). We tested each individual brain-*g* association, i.e. we fitted a separate SEM for each brain MRI measure. We then fitted a single SEM in which all indicators contributed to *g* variance; a so-called Multiple Indicators Multiple Causes (MIMIC; Muthen, 1989) model. We excluded TBV from this, to avoid model fit and theoretical part-whole issues. Significant correlated residual paths among the imaging variables estimated from modification indices were included. False-Discovery Rate (FDR; Benjamini and Hochberg, 1995) correction of the *p*-values (implemented using the p.adjust function in R) was applied across the six bivariate associations of interest, and then across each of the path estimates in the multivariate SEM. Manifest variables were corrected as described above.

We then conducted an additional—non-pre-registered—analysis, to investigate whether the substantially lower proportion of *g* variance accounted for by multiple global MRI measures in this sample, when compared to our prior work in an older cohort (Ritchie et al., 2015), was due to moderating effects of age. We split the sample by age into groups with equal-sized cognitive-MRI overlap (middle age N = 10,164; older age N = 10,166) to ensure no imbalance in statistical power (above and below age 63.29 years). Initially, we tested for measurement invariance of *g* between the two age groups. Specifically, we were interested in weak factorial invariance (equal factor loadings) rather than strong (equal loadings and intercepts; as defined by Widaman, Ferrer & Conger, 2010), given the expected difference in cognitive performance between middle and older participants. We did this by comparing two multi-group SEMs; in the first, the cognitive test loadings on *g* were freely estimated, whereas in the second they were constrained to be equal in the middle and older-aged groups. We used a chi-squared test, the Akaike Information Criterion (AIC), and the sample-adjusted Bayesian Information Criterion (saBIC), and an additional check of factor congruence (coefficient of factor congruence; Lorenzo-Seva & ten Berge, 2006) using the ‘*psych*’ package, to test whether there was a difference between these sub-models.

Congruence coefficients index the similarity between factor solutions; a congruence coefficient greater than .90 indicates an extremely high level of similarity of the factor solutions. Next, we used a different set of two multi-group models to test whether the associations between *g* and the MRI measures differed as a function of age. In the first, the *g*-brain associations were freely estimated, and in the second, they were constrained to be equal between the two age groups. All measures were corrected for sex, and the MRI measures were also corrected for MRI head position. The group *g* loadings were set to equality. We ran this test for both a single *g*-TBV association, and then where multiple global MRI measures (GM, NAWM, WMH, *g*FA, *g*MD) predicted *g*. Differences between the two multi-group SEMs were assessed with a chi-squared test, the AIC, and the saBIC.

#### 2.3.3 Sex differences in g-brain MRI associations

We then investigated sex differences in the size of the brain-cognitive associations. Before doing so, we tested for measurement invariance between the sexes by creating a multi-group SEM including just the cognitive tests, with sex as the grouping variable, and tested for strong measurement invariance (as defined by Widaman, Ferrer & Conger, 2010). If strong measurement invariance was found (i.e., the model with strong invariance does not fit significantly more poorly, by a chi-squared test, the AIC, and the saBIC, than one where factor loadings and intercepts are freely-estimated), we aimed to test a set of models where the brain-cognitive associations was fixed to equality across the two sub-models grouped by sex, and one where it was freely-estimated. We used a chi-squared test, the AIC, and the saBIC to test whether there was a difference between these sub-models (thus indicating that there is a sex difference). For these analyses, the variables were adjusted for all the above-mentioned covariates except sex.

#### 2.3.4 Associations between g and regional brain MRI measures

Finally, we examined associations between *g* and regional brain measures: i) the fractional anisotropy and mean diffusivity in 27 white matter pathways, ii) cortical volumes of 48 regions according to the Harvard-Oxford cortical atlas segmentations, and iii) 14 subcortical volumes (bilateral nucleus accumbens, amygdala, caudate, hippocampus, pallidum, putamen, thalamus). We applied FDR correction within each family of tests (across all cortical tests, and separately across the 27 tests of WM tracts for FA, and then for MD, and across all subcortical tests). We hypothesised that the associations between general intelligence and brain volumes across the cortex would be consistent with the Parieto-Frontal Integration Theory (P-FIT; Jung & Haier, 2007), and be strongest in lateral frontal, superior parietal and temporal regions. Likewise, we hypothesised that thalamic and association fibres, plus forceps minor will show the statistically largest associations with general intelligence. The additional use of subcortical volumes was an addition to our pre-registered plan; subcortical structure did not figure largely in the P-FIT (Jung & Haier., 2007), though more recent work has reported associations between intelligence and overall subcortical volume (Ritchie et al., 2015), caudate (Basten et al., 2015; Grazioplene et al., 2015; Rhein et al., 2014), hippocampal (Valdés Hernández et al., 2017) and thalamic volume (Bohlken et al., 2014).

## 3. Results

### 3.1 Estimating a latent general factor of general intelligence, ‘*g*’

Participant characteristics are shown in Table 1. The cognitive tests were all correlated with medium effect sizes according to Cohen (1992): the Pearson’s *r* range was |0.300 to 0.405|. A first principal component (without age and sex correction) accounted for 55% of the variance, with loadings ranging from |0.71 to 0.80| (Table S2). The confirmatory factor analysis, in which each indicator was not corrected for age and sex, had two fit indices (TLI and RMSEA) outside our pre-registered criteria (CFI = 0.973, TLI = 0.918, RMSEA = 0.078, SRMR = 0.024). Modification indices suggested the addition of a residual correlation between Verbal-Numerical Reasoning and Matrix Reasoning (*r* = 0.170), following which model fit was above our pre-registered threshold across all fit indices (CFI = 0.995, TLI = 0.969, RMSEA = 0.048, SRMR = 0.010). The general factor of cognitive ability accounted for 40% of the cognitive test score variance (standardised loadings were Matrix Reasoning = 0.550, Symbol-Digit = 0.626, Verbal-Numerical Reasoning = 0.532, Trail-Making part B = −0.794). When we corrected each cognitive test within the SEM for age and sex, keeping the abovementioned residual correlation, the model fit the data well (CFI = 0.998, TLI = 0.978, RMSEA = 0.030, SRMR = 0.004). The general factor of cognitive ability accounted for 32% of the cognitive test score variance (standardised loadings were Matrix Reasoning = 0.505, Symbol-Digit = 0.479, Verbal-Numerical Reasoning = 0.592, Trail-Making part B = −0.666).

### 3.2 Associations between general intelligence (*g*) and brain MRI measures

Results between *g* and global brain MRI measures are shown in Table 2 and Figure 3. SEM fit statistics are reported in Table S3, and residual correlations among the global brain tissue measures from the MIMIC model are reported in Table S4. In all models, the cognitive and MRI indicators were adjusted for age and sex, and the MRI indicators also adjusted for head positioning confounds. The latent factor of general intelligence was correlated with TBV at *r* = 0.275, *p* < 0.001. Associations with GM (*r* = 0.270) and NAWM (*r* = 0.254) were significantly larger than the other three tissue-specific measures (i.e., WMH, *g*FA, *g*MD; *p* for comparisons < 0.001). The associations of *g* with white matter hyperintensities, general fractional anisotropy, and general mean diffusivity had effect sizes (*r*) ranging from |0.060 to 0.122|.

**Table 2.** Associations between *g* and global MRI measures across the whole sample.

| Model | Individual |  |  | Simultaneous |  |  |
| --- | --- | --- | --- | --- | --- | --- |
| | Std. Est. | SE | $p$ | Std. Est. | SE | $p$ |
| TBV | 0.275 | 0.012 | <0.001 | - | - | - |
| GM | 0.270 | 0.012 | <0.001 | 0.181 | 0.019 | <0.001 |
| NAWM | 0.254 | 0.012 | <0.001 | 0.128 | 0.019 | <0.001 |
| WMH | -0.122 | 0.012 | <0.001 | -0.121 | 0.013 | <0.001 |
| $gFA$ | 0.084 | 0.011 | <0.001 | -0.014 | 0.017 | 0.495 |
| $gMD$ | -0.060 | 0.012 | <0.001 | -0.038 | 0.018 | 0.095 |

*Note.* Standardised estimates (Std. Est.) and standard errors (SE) reported. TBV: total brain volume, GM: grey matter volume, NAWM: normal-appearing white matter volume, WMH: white matter hyperintensity volume, $gFA$ : general factor of white matter fractional anisotropy, $gMD$ : general factor of white matter mean diffusivity. Manifest variables are corrected for age and sex; brain measures also corrected for head positioning confounds.

**Table 3.** Individual and unique contributions to *g* from global MRI measures across middle and older age groups.

| Model | Middle ($\leq 63.29$ yrs) | | | | Older ($> 63.29$ yrs) | | | |
| --- | --- | --- | --- | --- | --- | --- | --- | --- |
|  | Individual |  | Simultaneous |  | Individual |  | Simultaneous |  |
|  | *Std. Est.* | *p* | *Std. Est.* | *p* | *Std. Est.* | *p* | *Std. Est.* | *p* |
| TBV | 0.285 | <0.001 | - | - | 0.313 | <0.001 | - | - |
| *R*<sup>2</sup> | 0.081 |  | - | - | 0.098 |  | - | - |
| GM | 0.266<sup>a</sup> | <0.001 | 0.133 | <0.001 | 0.323<sup>a</sup> | <0.001 | 0.293 | <0.001 |
| NAWM | 0.275 | <0.001 | 0.165 | <0.001 | 0.275 | <0.001 | 0.066 | <0.001 |
| WMH | -0.129 | <0.001 | -0.114 | <0.001 | -0.174 | <0.001 | -0.149 | 0.007 |
| *g*FA | 0.106 | <0.001 | 0.032 | 0.097 | 0.109 | <0.001 | -0.089 | <0.001 |
| *g*MD | -0.060 |  | 0.001 | 0.585 | -0.121 |  | -0.115 | <0.001 |
| *R*<sup>2</sup> |  |  | 0.059 |  |  |  | 0.134 |  |

Note. Std. Est: standardised estimate. Groups split at 63.29 years. <sup>a</sup>Magnitudes were significantly different according to a $\chi^2$ test ( $p < 0.012$ , and significant following FDR). Models are corrected for sex; brain measures also corrected for head positioning confounds. All model fits were CFI and TLI $> 0.963$ , RMSEA and SRMR $< 0.022$ . Associations between *g* and TBV were not significantly different between middle and older ages: $\Delta\chi^2(1) = 0.784$ , $p = 0.376$ , $\Delta AIC = 1$ , $\Delta saBIC = 10$ . However, the magnitude of *g* associations with multiple global measures were significantly different between age groups: $\Delta\chi^2(7) = 188.02$ , $p = <0.001$ , $\Delta AIC = 174$ , $\Delta saBIC = 138$ . TBV: total brain volume, GM: grey matter volume, WM: white matter volume, WMH: white matter hyperintensity volume, FA: fractional anisotropy, MD: mean diffusivity.

**Figure 3.**
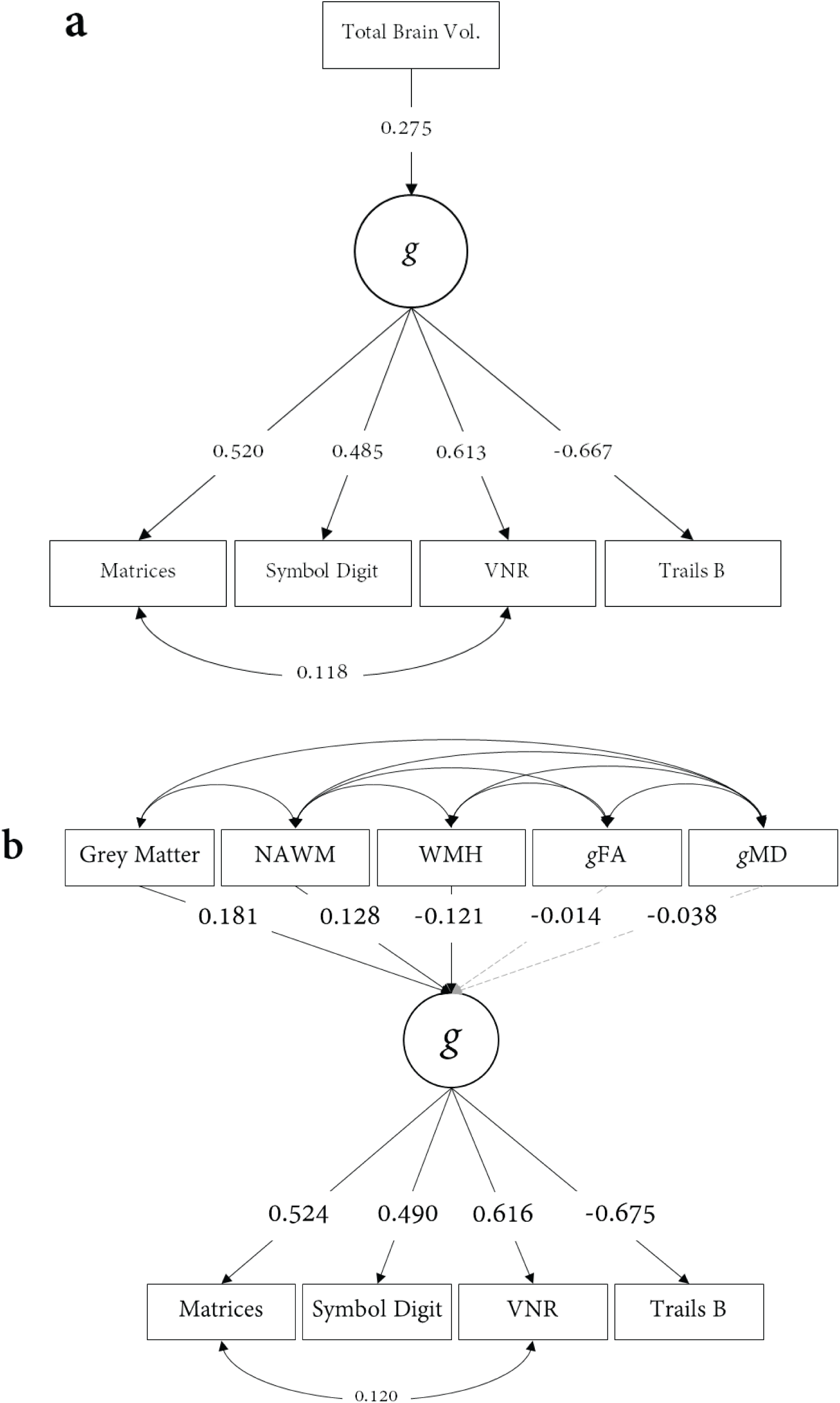
Associations between global brain MRI measures and *g*. Panel **a)** shows associations with total brain volume, and panel **b)** shows tissue-specific brain MRI measures accounting for 6.54% of the variance in *g*. Standardised estimates are reported; grey dashed paths are non-significant. Indicators are all corrected for age, sex, with imaging data also corrected for scanner head position coordinates. MRI residual correlations are shown in Table S5.

In a multivariate SEM (MIMIC model), we found that all MRI measures (GM, NAWM, WMH, *g*FA, *g*MD) accounted for 6.54% of the variance in *g*. The unique contributions to this variance were largest for grey matter volume (*β* = 0.181) with equivalent contributions of NAWM and WMH volumes (*β* = 0.128 and *β* = −0.121, respectively). Neither measure of white matter microstructure made significant unique contributions beyond this, following FDR correction (*g*FA *β* = −0.014, *p* = 0.403; and *g*MD *β* = −0.038, *p* = 0.035).

Ritchie et al. (2015) previously reported that the total variance in *g* explained by multiple MRI markers in an older cohort (all aged approximately 73 years) was 18-21%. To determine whether the difference between that estimate and the one reported here may be attributable to differences in the age range of the samples, we conducted a post-hoc supplementary test of differences in the proportion of variance explained by age. Initially, we tested whether *g* exhibited weak measurement invariance across the two age groups. Both models had excellent fit to the data, and were highly similar: saBIC indicated that weak invariance was preferred, contradicting the AIC results and the small but significant difference detected by the chi-squared test (Δχ^2^(3) = 27.617, *p* = <0.001, ΔAIC = 22, ΔsaBIC = −6.32). Comparing the magnitude and rank order of the freely estimated factor loadings between middle-aged (MR = 0.506, SDS = 0.539, VNR = 0.614, TMTb = −0.719) and older (MR = 0.522, SDS = 0.492, VNR = 0.569, TMTb = −0.701) participants also suggested that *g* exhibited weak factorial invariance between groups (coefficient of factor congruence = 1.00).

We then investigated whether the proportion of *g* variance accounted for by MRI measures was substantially greater in older than middle-aged participants. Results are shown in Table 3. A model with unconstrained *g-*MRI associations fitted the data significantly better than when the associations were constrained to equality between age groups (Δχ^2^(7) = 188.02, *p* = <0.001, ΔAIC = 174, ΔsaBIC = 138). In the model in which g-MRI associations were allowed to differ by age group, GM, NAWM, WMH, *g*FA and *g*MD together explained a total of 5.9% of the variance in *g* among the middle aged group, compared to 13.4% in older age. *g*-brain association magnitudes were all stronger in older age for GM (0.133 versus 0.293), WMH (−0.114 versus −0.149), *g*FA (0.032 versus −0.089) and *g*MD (0.001 versus −0.115); the one exception was NAWM volume (which showed stronger *g* associations in middle than older age; 0.165 versus 0.066). Notably, the global dMRI measures (*g*FA and *g*MD) were non-significant in the middle-age group. Moreover, we did not observe this significant age difference in variance explained in *g* by TBV alone, or by each individual MRI measure in isolation (all p-values for chi-squared tests were non-significant following FDR correction, with the exception of GM, which was more strongly associated with *g* in the older group, 0.266 versus 0.323; Δχ^2^(1) = 13.095, *p* < 0.001).

In summary, whereas the individual associations between brain measures and intelligence were of relatively similar magnitudes in middle and older age, when entered together they explained more than double the variance in *g* in older age. Inspection of the age differences in the correlational structure among these imaging markers (Figure S2) indicated that the source of increased variance explained is not likely to be attributable to their diverging collinearity (i.e. they overlap less in older age, and thus convey more unique information), given that the only notable differences were in associations between WMH and both gFA and gMD, which were stronger in older than younger age.

### 3.3 Sex differences in g-brain MRI associations

Before testing for sex differences in the size of the associations between *g* and brain MRI measures, we tested for measurement invariance between the sexes. We found that the model of strong factorial invariance of *g* did not fit more poorly than the model in which factor loadings and intercepts were freely-estimated (ΔAIC = 28, ΔsaBIC = −22, Δχ^2^ (4) = 40.027, *p* = <0.001). These results are reported in Table S5. We then tested for sex differences in the magnitude of associations between *g* and MRI measures. We did so by comparing two group models; one in which the brain-*g* association is fixed to equality between sexes, and the other in which it is freely estimated. We found that there were no significant sex differences in the magnitude of the association between *g* and any global brain MRI measure (*p* ≥ 0.171) except for NAWM (*p* = 0.011, standardised estimate for females = 0.185, males = 0.241), which was non-significant following FDR correction.

### 3.4 Associations between g and white matter microstructure

SEMs testing associations between *g* and the FA and MD of each white matter tract fitted the data well (all CFI ≥ 0.991, TLI ≥ 0.980, RMSEA ≤ 0.020, SRMR ≤ 0.016); results are shown in Figure 4, and Tables S6 and S7. Associations with *g* were in the expected direction, such that higher *g* was related to higher FA and lower MD (with the exception of the left medial lemniscus *β* = 0.027, FDR *q* = 0.018). Only a few pathways had non-significant associations with *g* (FA and MD in the bilateral acoustic radiations, FA in the middle cerebellar peduncle, and MD in the right parahippocampal cingulum, Forceps Major, bilateral inferior longitudinal fasciculus and left medial lemniscus). The effect sizes were not homogeneous across tracts (FA range = 0.001 to 0.112; MD range = −0.101 to 0.027). Consistent with our hypothesis, the magnitude of associations with *g* were numerically largest within thalamic pathways (FA mean = 0.080, MD mean = −0.081), and in association fibres and Forceps Minor (FA mean = 0.062, MD mean = −0.036) than within projection fibres and Forceps Major (FA mean = 0.033, MD mean = 0.009)^1^. However, it is also notable that both aspects of the cingulum bundle, showed among the weakest *g* relationship among association fibres, and that more generally there was a considerable amount of overlap between these classes of tract (for example, the right corticospinal tract MD was associated with *g* at levels comparable with most association fibres).

**Figure 4.**
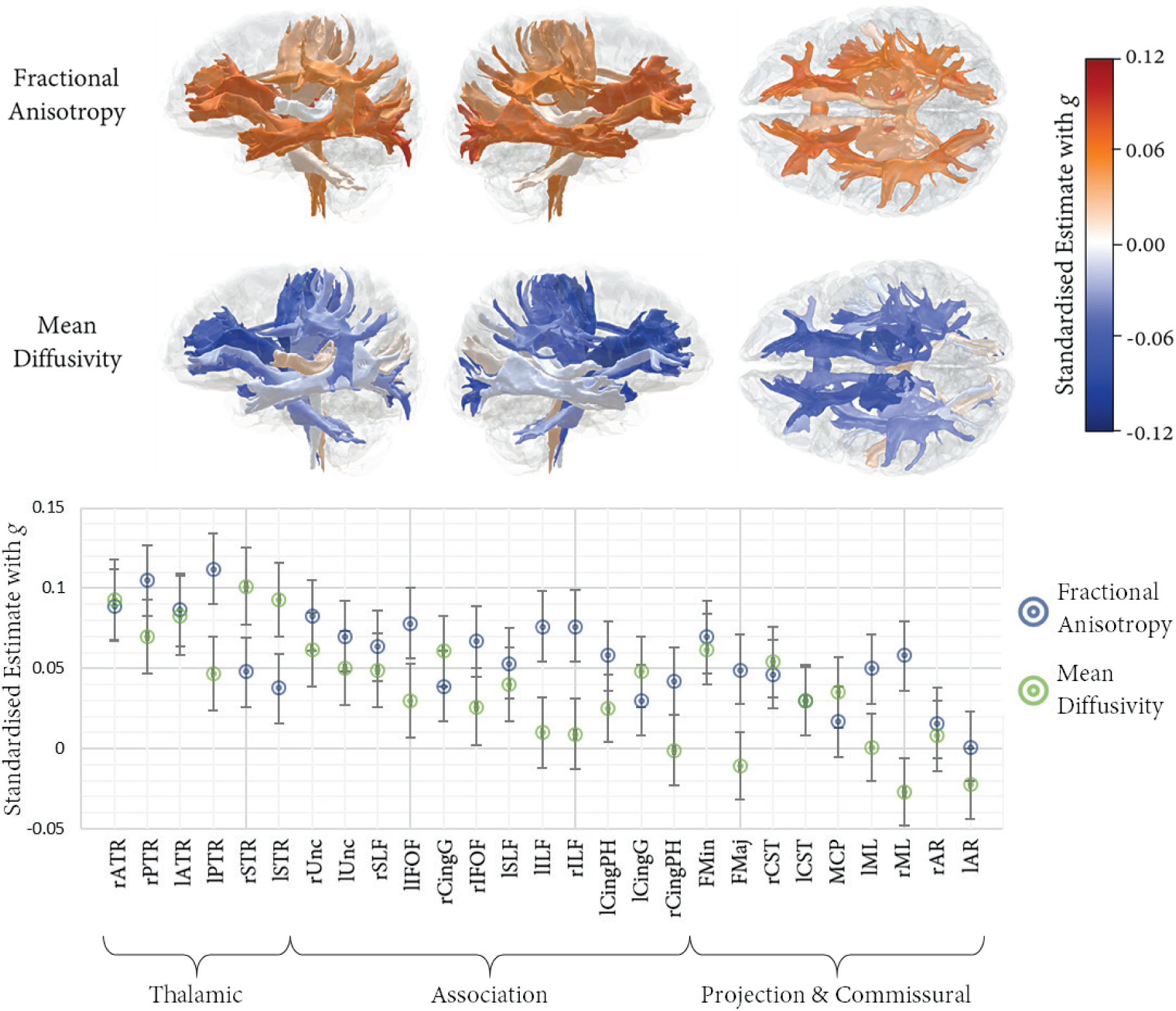
Associations between white matter tract-specific microstructure and *g* – MD valences have been flipped to aid visual comparison. Tabulated results also reported in Tables S3 and S4. Top panel shows left and right lateral and superior views of the white matter tracts of interest, heatmapped according to association magnitude. Lower panel displays the association magnitudes sorted by tract class and then from strongest to weakest (based on the average of FA and MD), with 95% CIs.

### 3.5 Associations between g and cortical regions

Associations between *g* and cortical regional volumes were all positive and all significant following FDR correction. The results are reported in Figure 5 and in Table S8; all models fitted the data well (CFI ≥ 0.990, TLI ≥ 0.979, RMSEA ≤ 0.021, SRMR ≤ 0.016). As with the white matter analyses above, there was regional heterogeneity in association magnitudes across the cortical surface. Substantial portions of the frontal lobe (frontal pole, frontal orbital, subcallosal) were among the numerically largest associations, bilaterally (range = 0.161 to 0.212), and these were significantly larger than other frontal regions (*p* < 0.001). Associations between the insula cortex and *g* (left = 0.186, right = 0.197) were also large compared to the average magnitude across all ROIs (M = 0.116, SD = 0.036). Associations within the temporal lobe were strongest in the temporal pole and anterior portions of the superior temporal gyrus (range = 0.140 to 0.160), and in more anterior portions of the parahippocampal and fusiform gyri (range = 0.148 to 0.154). Notably, the temporal lobe appeared to show a gradient of anterior > posterior for both lateral and medial portions, and the lateral surface also showed evidence of a superior > inferior gradient. Compared to the above-mentioned frontal, temporal and insula volumes, parietal regions were consistently and significantly more weakly associated with *g* (range = 0.066 to 0.101, *p* < 0.001). With the exception of the lingual, precuneus, and lateral occipital cortex (range = 0.109 to 0.150), occipital volumes were among the most weakly associated with *g* (range = 0.066 to 0.092).

**Figure 5.**
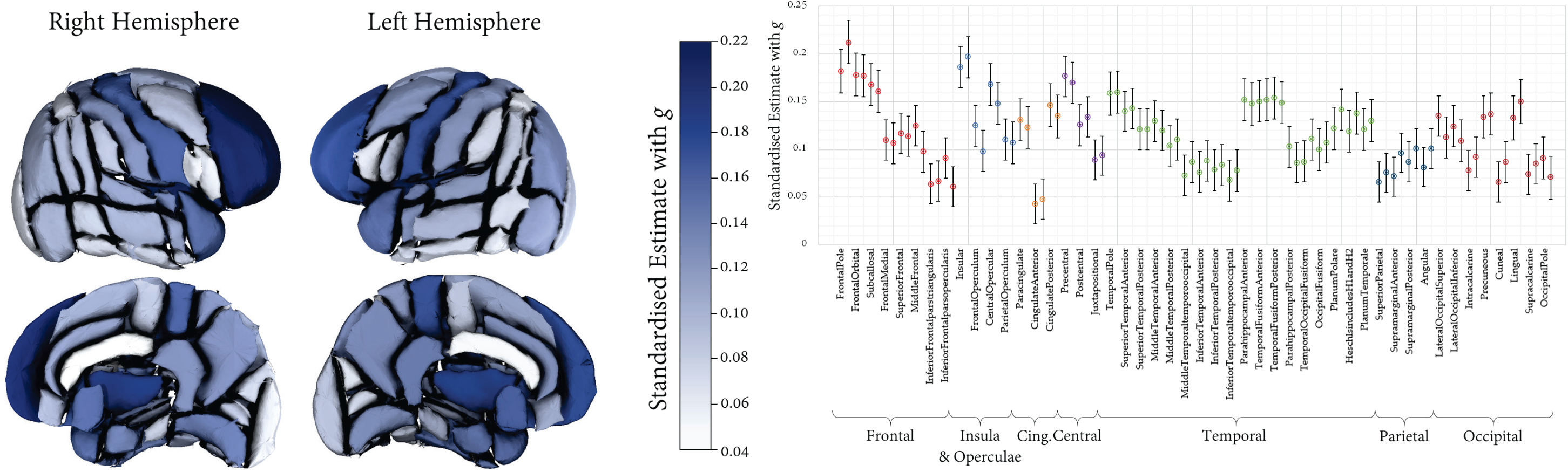
Associations between regional cortical volumes and *g* with 95% CIs. Left and right associations are shown separately (left hand regions appear first). Association magnitudes are also reported in Table S5.

### 3.6 Associations between g and subcortical volumes

As with the cortical analyses, subcortical volumes were all positively associated with *g*, and all were significant following FDR correction. The results are reported in Table S9. All models fitted the data well (CFI ≥ 0.990, TLI ≥ 0.979, RMSEA ≤ 0.020, SRMR ≤ 0.014), and standardised estimates ranged from 0.050 to 0.277. Largest effect sizes were found for the Thalamus (left = 0.274, right = 0.277), which was significantly larger than for all other subcortical regions (*p* < 0.001). The Amygdala showed the weakest associations of all (left = 0.063, right = 0.050), whereas the remaining volumes (Accumbens, Caudate, Hippocampus, Pallidum and Putamen) showed comparable magnitudes (range = 0.090 to 0.146).

## Discussion

In this sample of 29,004 middle and older aged participants, we found that the association between total brain volume and a latent factor of general intelligence was *r* = 0.275. The current single-cohort analysis was not affected by the confounds of cross-cohort heterogeneity in the protocol for intelligence and brain size measurement which have affected recent meta-analyses of this association (McDaniel, 2005; Rushton & Ankney, 2009; Pietschnig et al., 2015; Gignac & Bates, 2017. This estimate is equal to the mid-point between the meta-analytic effect size estimate from Pietschnig et al., (2015; *r* = 0.24) and the quality-corrected estimate of Gignac and Bates (2017; *r* = 0.31). It is also considerably larger than previous estimates using a single cognitive indicator (verbal numerical reasoning) in an earlier UK Biobank release (*r* = 0.19, N = 13,608; Nave et al., 2018; *r* = 0.177, N = 5,216; Ritchie et al., 2018), emphasising the utility of our latent variable approach, which was also informed by a larger sample. The fact that the association between *g* and TBV was not significantly different between sexes is in contrast the results reported by McDaniel (2005), but not with a larger, more recent meta-analysis (Pietschnig et al., 2015), and prior work in an earlier UK Biobank release using the VNR only (Ritchie et al., 2018), adding weight to the hypothesis that more specific brain characteristics compensate for the large brain size difference between males and females.

We also ascertained that the *g*-TBV association belies important heterogeneity at both the global tissue level, and at the regional level across cortex, subcortex and white matter. GM and NAWM were the strongest global tissue correlates of *g*, with WMH and microstructural measures showing weaker but significant associations in separate models. However, when modelled simultaneously in a MIMIC model, unique contributions of WMH and NAWM were near identical (GM still largest), whereas information about white matter microstructure apparently not carrying any unique information about individual differences in *g*. Together, these measures explained only 6.54% of *g* variance. This was substantially lower than prior estimates using similar global structural brain metrics (Ritchie et al., 2015) which explained as much as 21% *g* variance. Given that their participants were from an older cohort – all participants were born in 1936 and were approximately 73 years old at scanning – we conducted a post-hoc analysis to ascertain whether the brain measures would account for more variance in older than in younger participants. We found that these measures accounted for more than double the *g* variance in older participants compared to those in middle-age (13.4% and 5.9%, respectively); thus while still smaller than that accounted for in Ritchie et al (2015), this does support the notion that age moderates the relationship between general intelligence and multiple aspects of brain tissue structure. This age moderation pattern was only observed in a multi-predictor analysis that simultaneously included multiple MRI-based predictors and not in an analysis of total brain volume or individual MRI predictor alone.

First, these age differences stood in contrast to the apparent age invariance of the association between *g* and total brain volume (as found by Pietschnig et al., 2015). This supports the notion that total brain volume is a proxy for several other aspects of brain integrity whose variances are i) uniquely informative for cognitive function and ii) more informative than brain size alone. Moreover, total brain volume is likely to be an age-varying indicator of brain integrity, raising questions about the value of considering the brain size-intelligence relationship, in isolation, for furthering our mechanistic insight into the cerebral basis of intelligence. Overall, the age moderation pattern suggests that *g* may be less sensitive to the variance around ‘healthier’ / ‘younger’ averages (higher GM and NAWM volume, lower WMH volume, higher FA and lower MD).

In our analysis of regional brain correlates of intelligence, the cortical and grey matter associations were stronger than for the regional white matter microstructural parameters, on average, though association magnitudes were all of small effect size (Cohen, 1992). Nearly all regional measures were significant following FDR correction, and our findings of regional heterogeneity of *g* associations across the tissues of the brain were partly consistent with our hypotheses based on the P-FIT (Jung & Haier, 2007). Specifically, we hypothesised that *g* would be most strongly positively associated with lateral frontal, superior parietal and temporal cortical volumes, and show stronger (positive for FA, negative for MD) relationships with *g* in thalamic and association fibres, plus forceps minor. In accordance with this, we found relatively stronger associations in regions such as the frontal pole, dorsolateral frontal cortex, paracingulate, anterior aspects of both lateral and medial temporal lobes, and lateral occipital cortex. However, the comparatively weaker associations in inferior frontal, anterior cingulate, and superior parietal / angular / supramarginal areas were less consistent with P-FIT. Moreover, medial frontal regions (orbitofrontal and subcallosal), central and precentral gyri were among the strongest associations here, but were not explicitly implicated in prior reviews (Jung & Haier, 2007; Basten et al., 2015), and we also found associations with the insula and precuneus / posterior cingulate volumes which were only more recently implicated in general intelligence (Basten et al., 2015), and concurs with more recent insights into the dense and wide-ranging connectivity profile of the insula (Nomi et al., 2018). With reference to the white matter pathways, magnitudes were consistently smaller than for cortical regional volumes, but were strongest among thalamic and most association pathways, along with the forceps minor, which facilitate connectivity across many of the distal cortical regions highlighted by the P-FIT model.

Finally, we opted to include subcortical volumes in a post-hoc (non-pre-registered) analysis. Consistent with prior reports (Basten et al., 2015; Grazioplene et al., 2015; Rhein et al., 2014), we found significant bilateral associations with the caudate, though these were not significantly larger than the magnitudes found for the majority of subcortical structures. In fact, thalamic volume was substantially more strongly related to general intelligence (≥1.5 times as large) than any other subcortical structure (*r* for left and right *=* 0.255 and 0.251). This finding is in line with the highly complex connectivity profile of the thalamus, whose various nuclei share connections across much of the cortex (including prefrontal and hippocampal pathways; Behrens et al., 2003; Aggleton et al., 2010), its role in orchestrating cortical activity as well as an information relay (Rikhye et al., 2018), and a prior report of its phenotypic and genetic associations with intelligence (Bohlken et al., 2014). It is also consistent with previously-reported associations of the thalamus and its radiations with ageing, and to potential determinants thereof, such as vascular risk (Cox et al., 2016; Cox et al., 2019). However, it is also notable that the association between *g* and all subcortical structures, though not as large as for the Thalamus, were still comparable or larger than those exhibited by white matter microstructural measures.

The study has several limitations. The information reported here is correlational in nature, and though it describes what intelligent brains look like (insofar as these are some of the axes along which brains differ as a function of intelligence), it cannot directly differentiate between regions that are and are not *required* to support the cognitive processes subsumed beneath the umbrella of *g*. Neverthelss, it continues to be of interest and value to robustly quantify how and where brain structure and intelligence are associated; along with longitudinal data, lesion studies and other methodologies, such studies will help to triangulate the contributions that brain regions play in giving rise to individual differences in *g*. The study sample is also range restricted in three respects. First, they are members of a voluntary research study (and UK Biobank is known to be range restricted in some ways compared to the general population; Fry et al., 2017), Second, we know that the brain imaging subset of UK Biobank participants tends to live in less deprived areas (Lyall et al., 2019), and third, the age range of the sample is restricted to middle and older age. These sample limitations may affect the generalisability of our results with respect to the total brain, global tissue or regional results, and the pattern of age moderation; these whole-life-course patterns could more optimally be addressed in a large scale multi-cohort mega-analytic framework. Though the reliability of the more recent cognitive tests that we used here is not known, we note that the loadings and proportion of variance explained would be very unlikely to occur if they were unreliable tests, and that these compare favourably with the correlational structure of other UK Biobank cognitive tests where test-retest reliability is known to be low (Lyall et al., 2014). Moreover, the tests selected here were based on well-validated cognitive tests, and a paper covering their design and reporting results of a validation study of this enhanced cognitive battery in UK Biobank is the subject of ongoing work by the authors (CFR and IJD). Finally, it could be argued that the brain imaging methods might limit the fidelity with which we can measure the regional specificity of *g* associations across the brain. The 27 major pathways have the advantage of being well characterised and aid consistent identification across subjects, but they do not allow a direct measure of the WM connectivity between two cortical or subcortical sites of the brain, which would allow for a more precise and stringent test of *g* associations with the WM pathways underlying the P-FIT, as well as a less biased set of pathways (for example, the current dataset has more information on thalamic connectivity than on other subcortical pathways). Similarly, the cortical parcellation used here was one of convenience and does not correspond directly onto the Brodmann Areas used by Jung and Haier (2007), which makes mapping the current findings onto prior hypotheses opaque. For example, whereas the Harvard-Oxford atlas includes a paracingulate region (Brodmann Area 32), this additional cortical fold is not always present, and thus is perhaps more usefully referred to as “superior medial” cortex (e.g. see Cox et al., 2014). Likewise, frontal pole region subsumes a large portion of the frontal lobe compared to that described by Brodmann; though this concordance issue is well-known, and there is no straightforward solution (Bohland et al., 2009; Cox et al., 2014), it is important to interpret the results with these limitations in mind.

In conclusion, this preregistered study provides a large single sample analysis of the global and regional brain correlates of a latent factor of general intelligence. Our study design avoids issues of publication bias and inconsistent cognitive measurement to which meta-analyses are susceptible, and also provides a latent measure of intelligence which compares favourably with previous single-indicator studies of this type. We estimate the correlation between total brain volume and intelligence to be *r* = 0.275, which applies to both males and females. Multiple global tissue measures account for around double the variance in *g* in older participants, relative to those in middle age. Finally, we find that associations with intelligence were strongest in frontal, insula, anterior and medial temporal, lateral occipital and paracingulate cortices, alongside subcortical volumes (especially the thalamus) and the microstructure of the thalamic radiations, association pathways and forceps minor.

## Acknowledgements

We thank the UK Biobank participants and the UK Biobank team for their work in collecting, processing and disseminating these data for analysis. We also thank David C. Liewald for data and technical computing support. This research was conducted, using the UK Biobank Resource under approved project 10279, in The University of Edinburgh Centre for Cognitive Ageing and Cognitive Epidemiology (CCACE) (http://www.ccace.ed.ac.uk), part of the cross-council Lifelong Health and Wellbeing Initiative (MR/K026992/1). Funding from the Biotechnology and Biological Sciences Research Council (BBSRC) and Medical Research Council (MRC) is gratefully acknowledged. SRC and IJD were supported by MRC grants MR/M013111/1 and MR/R024065/1. IJD is additionally supported by the Dementias Platform UK (MR/L015382/1), and he, SRC and SJR by the Age UK-funded Disconnected Mind project (http://www.disconnectedmind.ed.ac.uk). SRC, SJR, IJD and EMT-D were supported by a National Institutes of Health (NIH) research grant R01AG054628. EMT-D was also supported by National Institutes of Health (NIH) research grant R01HD083613, and is a member of the Population Research Center at the University of Texas at Austin, which is supported by NIH center grant P2CHD042849. CF-R is supported by Dementias Platform UK (DPUK), funded through the MRC (MR/L023784/2).

## Supplementary Material

**Table S1.** Self-reported health variables for exclusion criteria

| Variable | Condition | Field ID | Code |
| --- | --- | --- | --- |
| *Health* |  |  |  |
|  | Illness code | 20002 |  |
|  | Dementia or Alzheimer's disease |  | 1263 |
|  | Parkinson's disease |  | 1262 |
|  | Chronic degenerative neurological |  | 1258 |
|  | Guillain-Barré syndrome |  | 1256 |
|  | Multiple Sclerosis |  | 1261 |
|  | Other demyelinating disease |  | 1397 |
|  | Stroke or ischaemic stroke |  | 1081 |
|  | Brain cancer |  | 1032 |
|  | Brain haemorrhage |  | 1491 |
|  | Brain/intracranial abscess |  | 1245 |
|  | Cerebral aneurysm |  | 1425 |
|  | Cerebral palsy |  | 1433 |
|  | Encephalitis |  | 1246 |
|  | Epilepsy |  | 1264 |
|  | Head injury |  | 1266 |
|  | Infections of the nervous system |  | 1244 |
|  | Ischaemic stroke |  | 1583 |
|  | Meningeal cancer |  | 1031 |
|  | Meningioma (benign) |  | 1659 |
|  | Meningitis |  | 1247 |
|  | Motor Neuron Disease |  | 1259 |
|  | Neurological injury/trauma |  | 1240 |
|  | Spina bifida |  | 1524 |
|  | Subdural haematoma |  | 1083 |
|  | Subarachnoid haemorrhage |  | 1086 |
|  | Transient ischaemic attack |  | 1082 |
| *Cognitive* |  |  |  |
|  | Symbol-Digit | 23324 |  |
|  | Matrix Reasoning | 6373 |  |
|  | VNR | 20016 |  |
|  | TMTb | 6350 |  |
| *Neuroimaging* |  |  |  |
|  | TBV | 25010 |  |
|  | GM | 25006 |  |
|  | WM | 25008 |  |
|  | WMH | 25781 |  |
|  | Cortical Volumes | 25782:25877 |  |
|  | Subcortical Volumes | 25011:25024 |  |
|  | WM Tract FA | 25488:25514 |  |
|  | WM Tract MD | 25515:25541 |  |

*Note.* VNR: verbal numerical reasoning, TMTb: Trail Making Test Part B, TBV: total brain volume, GM: grey matter volume, WM: white matter volume, WMH: white matter hyperintensity volume, FA: fractional anisotropy, MD: mean diffusivity.

**Table S2.** Principal components analysis of the four cognitive tests.

|  | PC1 | PC2 | PC3 | MR | DSS | VNR | TMTb |
| --- | --- | --- | --- | --- | --- | --- | --- |
| **Matrix Reasoning** | 0.73 | 0.27 | 0.62 | ***** | 0.360 | 0.405 | -0.302 |
| **Symbol Digit** | 0.72 | -0.57 | -0.01 | <0.001 | ***** | 0.301 | -0.343 |
| **VNR** | 0.71 | 0.53 | -0.41 | <0.001 | <0.001 | ***** | -0.300 |
| **TMTb** | -0.80 | 0.20 | 0.20 | <0.001 | <0.001 | <0.001 | ***** |
| **ProportionVar** | 0.55 | 0.18 | 0.15 | - | - | - | - |
| **CumulativeVar** | 0.55 | 0.73 | 0.88 | - | - | - | - |

Note. Loadings of the cognitive tests on the components (left three columns) and correlations among the cognitive tests (Pearson's $r$ ; right four columns) are shown. VNR: verbal numerical reasoning, TMTb: Trail Making Test Part B.

**Table S3.**
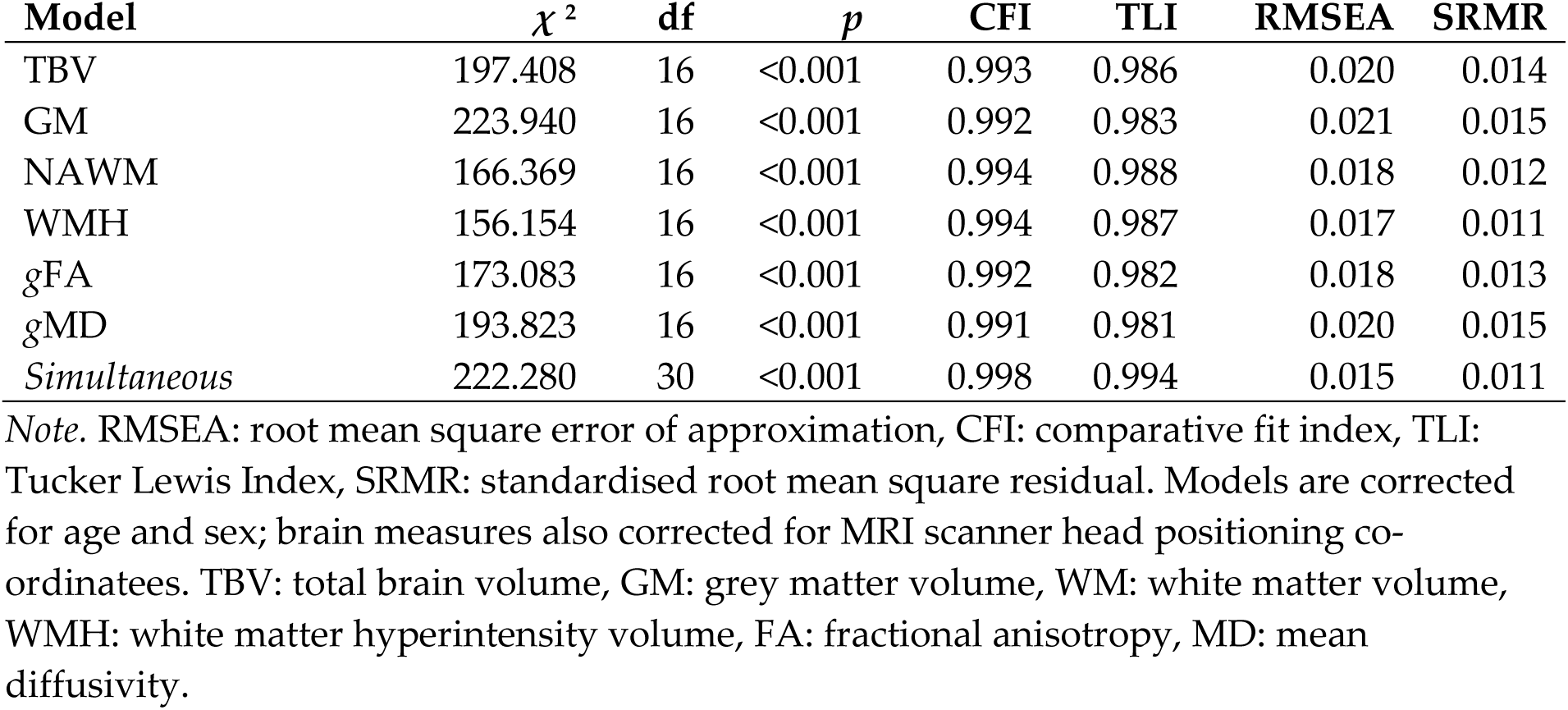
Structural equation model fit statistics for *g* with global MRI measures.

**Table S4.**
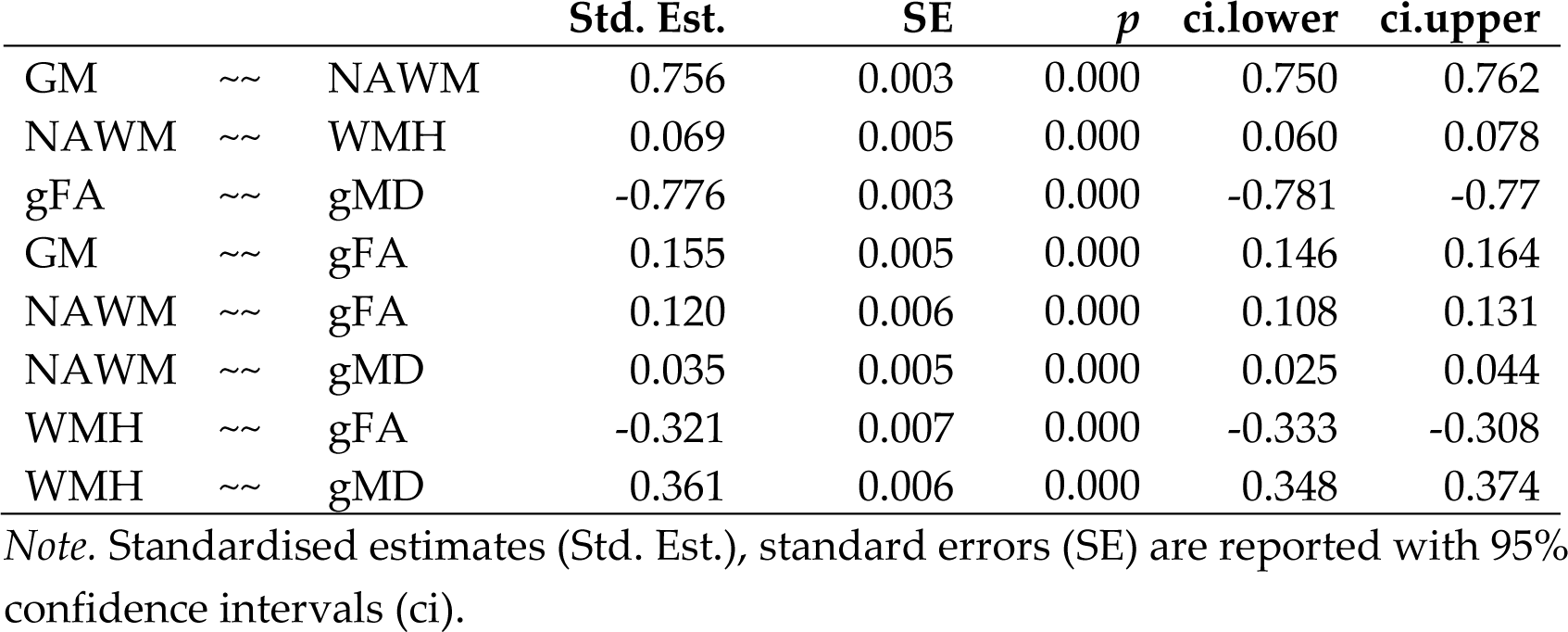
Residual correlations among global brain imaging measures from SEM Figure 3b.

| | | | Std. Est. | SE | $p$ | ci.lower | ci.upper |
| --- | --- | --- | --- | --- | --- | --- | --- |
| GM | ~~ | NAWM | 0.756 | 0.003 | 0.000 | 0.750 | 0.762 |
| NAWM | ~~ | WMH | 0.069 | 0.005 | 0.000 | 0.060 | 0.078 |
| gFA | ~~ | gMD | -0.776 | 0.003 | 0.000 | -0.781 | -0.77 |
| GM | ~~ | gFA | 0.155 | 0.005 | 0.000 | 0.146 | 0.164 |
| NAWM | ~~ | gFA | 0.120 | 0.006 | 0.000 | 0.108 | 0.131 |
| NAWM | ~~ | gMD | 0.035 | 0.005 | 0.000 | 0.025 | 0.044 |
| WMH | ~~ | gFA | -0.321 | 0.007 | 0.000 | -0.333 | -0.308 |
| WMH | ~~ | gMD | 0.361 | 0.006 | 0.000 | 0.348 | 0.374 |

Note. Standardised estimates (Std. Est.), standard errors (SE) are reported with 95% confidence intervals (ci).

**Table S5.** Structural equation model estimates and fit statistics testing factorial invariance of *g* between males and females.

| Model<br>Parameters | Baseline |  | Strong Invariance |  | Model<br>Comparison |
| --- | --- | --- | --- | --- | --- |
|  | Females | Males | Females | Males |  |
| g Loadings |  |  |  |  |  |
| Matrix Reasoning | 1.000 | 1.000 | 1.000 | 1.000 |  |
| Symbol-Digit | 2.335 | 2.372 | 2.356 | 2.356 |  |
| VNR | 1.126 | 1.149 | 1.139 | 1.139 |  |
| Trail Making B | -0.214 | -0.220 | -0.218 | -0.218 |  |
| Matrix Reasoning<br>~~ VNR | 0.511 | 0.232 | 0.522 | 0.226 |  |
| Intercepts |  |  |  |  |  |
| Matrix Reasoning | 12.352 | 12.983 | 12.614 | 12.614 |  |
| Symbol Digit | 39.294 | 38.757 | 38.842 | 38.842 |  |
| VNR | 8.147 | 8.554 | 8.273 | 8.273 |  |
| Trail Making B | 5.051 | 5.085 | 5.083 | 5.083 |  |
| Model Fit |  |  |  |  |  |
| CFI |  | 0.998 |  | 0.998 |  |
| TLI |  | 0.984 |  | 0.995 |  |
| RMSEA |  | 0.030 |  | 0.017 |  |
| SRMR |  | 0.005 |  | 0.008 |  |
| AIC |  | 272370 |  | 272370 | 0 |
| saBIC |  | 272544 |  | 272513 | -31 |
| $\chi^2$ | | 28.445 | | 40.266 | 11.821 |
| (df) |  | 2 |  | 8 | 6 |
| p |  | <0.001 |  | <0.001 | 0.066 |

*Note.* Unstandardised estimates are reported. VNR: verbal numerical reasoning, CFI: comparative fit index, TLI: Tucker Lewis Index, RMSEA: root mean square error of approximation, SRMR: standardised root mean square residual, AIC: Akaike Information Criterion, saBIC: sample-adjusted Bayesian Information Criterion. Models are corrected for age.

**Table S6.** Associations between *g* and white matter tract fractional anisotropy.

| | Estimate | SE | $p$ | ci.lower | ci.upper | FDR $q$ |
| --- | --- | --- | --- | --- | --- | --- |
| lAR | 0.001 | 0.011 | 0.896 | -0.020 | 0.023 | 0.896 |
| rAR | 0.016 | 0.011 | 0.150 | -0.006 | 0.038 | 0.156 |
| lATR | 0.087 | 0.011 | 0.000 | 0.064 | 0.109 | 0.000 |
| rATR | 0.089 | 0.011 | 0.000 | 0.067 | 0.112 | 0.000 |
| lCingG | 0.030 | 0.011 | 0.007 | 0.008 | 0.052 | 0.008 |
| rCingG | 0.039 | 0.011 | 0.001 | 0.017 | 0.061 | 0.001 |
| lCingPH | 0.058 | 0.011 | 0.000 | 0.036 | 0.079 | 0.000 |
| rCingPH | 0.042 | 0.011 | 0.000 | 0.021 | 0.063 | 0.000 |
| lCST | 0.030 | 0.011 | 0.008 | 0.008 | 0.051 | 0.009 |
| rCST | 0.046 | 0.011 | 0.000 | 0.025 | 0.068 | 0.000 |
| FMaj | 0.049 | 0.011 | 0.000 | 0.028 | 0.071 | 0.000 |
| FMin | 0.070 | 0.012 | 0.000 | 0.047 | 0.092 | 0.000 |
| lIFOF | 0.078 | 0.011 | 0.000 | 0.056 | 0.100 | 0.000 |
| rIFOF | 0.067 | 0.011 | 0.000 | 0.045 | 0.089 | 0.000 |
| lILF | 0.076 | 0.011 | 0.000 | 0.054 | 0.098 | 0.000 |
| rILF | 0.076 | 0.011 | 0.000 | 0.054 | 0.099 | 0.000 |
| MCP | 0.017 | 0.011 | 0.135 | -0.005 | 0.039 | 0.145 |
| lML | 0.050 | 0.011 | 0.000 | 0.028 | 0.071 | 0.000 |
| rML | 0.058 | 0.011 | 0.000 | 0.036 | 0.079 | 0.000 |
| lPTR | 0.112 | 0.011 | 0.000 | 0.090 | 0.134 | 0.000 |
| rPTR | 0.105 | 0.011 | 0.000 | 0.083 | 0.127 | 0.000 |
| lSLF | 0.053 | 0.011 | 0.000 | 0.031 | 0.075 | 0.000 |
| rSLF | 0.064 | 0.011 | 0.000 | 0.042 | 0.086 | 0.000 |
| lSTR | 0.038 | 0.011 | 0.001 | 0.016 | 0.059 | 0.001 |
| rSTR | 0.048 | 0.011 | 0.000 | 0.026 | 0.069 | 0.000 |
| lUnc | 0.070 | 0.011 | 0.000 | 0.048 | 0.092 | 0.000 |
| rUnc | 0.083 | 0.011 | 0.000 | 0.061 | 0.105 | 0.000 |

*Note.* Standardised estimates, standard errors (SE) are reported. Right and left are denoted by l and r. AR: acoustic radiation, ATR: anterior thalamic radiation, Cing: cingulum bundle gyrus (G) and parahippocampal (PH), CST: corticospinal tract, FMaj: forceps major, FMin: forceps minor, IFOF: inferior fronto-occipital fasciculus, ILF: inferior longitudinal fasciculus, MCP: middle cerebellar peduncle, ML: medial lemniscus, PTR: posterior thalamic radiation, SLF: superior longitudinal fasciculus, STR: superior thalamic radiation, Unc: uncinate. All model fits were $\chi^2(17) \leq 285.419$ $p < 0.001$ , CFI $\geq 0.985$ , TLI $\geq 0.970$ , RMSEA $\leq 0.023$ , SRMR $\leq 0.015$ .

**Table S7.** Associations between *g* and white matter tract mean diffusivity.

| | Estimate | SE | $p$ | ci.lower | ci.upper | FDR $q$ |
| --- | --- | --- | --- | --- | --- | --- |
| lAR | 0.022 | 0.011 | 0.042 | 0.001 | 0.044 | 0.053 |
| rAR | -0.008 | 0.011 | 0.486 | -0.029 | 0.014 | 0.525 |
| lATR | -0.083 | 0.013 | 0.000 | -0.108 | -0.058 | 0.000 |
| rATR | -0.093 | 0.013 | 0.000 | -0.118 | -0.068 | 0.000 |
| lCingG | -0.048 | 0.011 | 0.000 | -0.071 | -0.026 | 0.000 |
| rCingG | -0.061 | 0.011 | 0.000 | -0.083 | -0.039 | 0.000 |
| lCingPH | -0.025 | 0.011 | 0.022 | -0.047 | -0.004 | 0.031 |
| rCingPH | 0.001 | 0.011 | 0.919 | -0.020 | 0.023 | 0.919 |
| lCST | -0.030 | 0.011 | 0.007 | -0.052 | -0.008 | 0.012 |
| rCST | -0.054 | 0.011 | 0.000 | -0.076 | -0.032 | 0.000 |
| FMaj | 0.011 | 0.011 | 0.320 | -0.011 | 0.032 | 0.393 |
| FMin | -0.062 | 0.012 | 0.000 | -0.085 | -0.040 | 0.000 |
| lIFOF | -0.030 | 0.012 | 0.012 | -0.052 | -0.007 | 0.018 |
| rIFOF | -0.026 | 0.012 | 0.031 | -0.049 | -0.002 | 0.042 |
| lILF | -0.010 | 0.011 | 0.370 | -0.033 | 0.012 | 0.434 |
| rILF | -0.009 | 0.012 | 0.427 | -0.032 | 0.013 | 0.480 |
| MCP | -0.035 | 0.011 | 0.002 | -0.057 | -0.013 | 0.003 |
| lML | -0.001 | 0.011 | 0.894 | -0.023 | 0.020 | 0.919 |
| rML | 0.027 | 0.011 | 0.012 | 0.006 | 0.048 | 0.018 |
| lPTR | -0.047 | 0.012 | 0.000 | -0.070 | -0.024 | 0.000 |
| rPTR | -0.070 | 0.012 | 0.000 | -0.093 | -0.047 | 0.000 |
| lSLF | -0.040 | 0.012 | 0.001 | -0.063 | -0.017 | 0.001 |
| rSLF | -0.049 | 0.012 | 0.000 | -0.072 | -0.026 | 0.000 |
| lSTR | -0.093 | 0.012 | 0.000 | -0.117 | -0.070 | 0.000 |
| rSTR | -0.101 | 0.012 | 0.000 | -0.125 | -0.077 | 0.000 |
| lUnc | -0.050 | 0.012 | 0.000 | -0.073 | -0.027 | 0.000 |
| rUnc | -0.062 | 0.012 | 0.000 | -0.085 | -0.039 | 0.000 |

*Note.* Standardised estimates, standard errors (SE) are reported. AR: acoustic radiation, ATR: anterior thalamic radiation, Cing: cingulum bundle gyrus (G) and parahippocampal (PH), CST: corticospinal tract, FMaj: forceps major, FMin: forceps minor, IFOF: inferior fronto-occipital fasciculus, ILF: inferior longitudinal fasciculus, MCP: middle cerebellar peduncle, ML: medial lemniscus, PTR: posterior thalamic radiation, SLF: superior longitudinal fasciculus, STR: superior thalamic radiation, Unc: uncinate. All model fits were $\chi^2(17) \leq 303.473$ $p < 0.001$ , CFI $\geq 0.985$ , TLI $\geq 0.970$ , RMSEA $\leq 0.024$ , SRMR $\leq 0.017$ .

**Table S8.** Associations between *g* and grey matter regional volumes.

|  | Estimate | SE | *p* | ci.lower | ci.upper |
| --- | --- | --- | --- | --- | --- |
| FrontalPoleleft | 0.182 | 0.012 | 0.000 | 0.159 | 0.205 |
| FrontalPoleright | 0.212 | 0.011 | 0.000 | 0.189 | 0.234 |
| InsularCortexleft | 0.186 | 0.011 | 0.000 | 0.164 | 0.207 |
| InsularCortexright | 0.197 | 0.011 | 0.000 | 0.176 | 0.219 |
| SuperiorFrontalGyrusleft | 0.117 | 0.011 | 0.000 | 0.096 | 0.138 |
| SuperiorFrontalGyrusright | 0.114 | 0.011 | 0.000 | 0.093 | 0.135 |
| MiddleFrontalGyrusleft | 0.125 | 0.011 | 0.000 | 0.104 | 0.146 |
| MiddleFrontalGyrusright | 0.098 | 0.011 | 0.000 | 0.077 | 0.120 |
| InferiorFrontalGyrusparstriangularisleft | 0.064 | 0.011 | 0.000 | 0.043 | 0.085 |
| InferiorFrontalGyrusparstriangularisright | 0.067 | 0.011 | 0.000 | 0.046 | 0.088 |
| InferiorFrontalGyrusparsopercularisleft | 0.091 | 0.011 | 0.000 | 0.070 | 0.112 |
| InferiorFrontalGyrusparsopercularisright | 0.061 | 0.011 | 0.000 | 0.040 | 0.082 |
| PrecentralGyrusleft | 0.177 | 0.011 | 0.000 | 0.156 | 0.199 |
| PrecentralGyrusright | 0.170 | 0.011 | 0.000 | 0.149 | 0.192 |
| TemporalPoleleft | 0.159 | 0.011 | 0.000 | 0.137 | 0.182 |
| TemporalPoleright | 0.160 | 0.011 | 0.000 | 0.138 | 0.182 |
| SuperiorTemporalGyrusanteriorleft | 0.140 | 0.011 | 0.000 | 0.119 | 0.161 |
| SuperiorTemporalGyrusanteriorright | 0.143 | 0.011 | 0.000 | 0.122 | 0.164 |
| SuperiorTemporalGyrusposteriorleft | 0.121 | 0.011 | 0.000 | 0.099 | 0.142 |
| SuperiorTemporalGyrusposteriorright | 0.121 | 0.011 | 0.000 | 0.099 | 0.142 |
| MiddleTemporalGyrusanteriorleft | 0.130 | 0.011 | 0.000 | 0.109 | 0.152 |
| MiddleTemporalGyrusanteriorright | 0.120 | 0.011 | 0.000 | 0.099 | 0.141 |
| MiddleTemporalGyrusposteriorleft | 0.104 | 0.011 | 0.000 | 0.083 | 0.126 |
| MiddleTemporalGyrusposteriorright | 0.110 | 0.011 | 0.000 | 0.088 | 0.131 |
| MiddleTemporalGyrustemporooccipitalleft | 0.073 | 0.011 | 0.000 | 0.052 | 0.094 |
| MiddleTemporalGyrustemporooccipitalright | 0.087 | 0.011 | 0.000 | 0.066 | 0.109 |
| InferiorTemporalGyrusanteriorleft | 0.076 | 0.011 | 0.000 | 0.054 | 0.097 |
| InferiorTemporalGyrusanteriorright | 0.088 | 0.011 | 0.000 | 0.067 | 0.109 |
| InferiorTemporalGyrusposteriorleft | 0.079 | 0.011 | 0.000 | 0.058 | 0.101 |
| InferiorTemporalGyrusposteriorright | 0.084 | 0.011 | 0.000 | 0.063 | 0.106 |
| InferiorTemporalGyrustemporooccipitalleft | 0.068 | 0.011 | 0.000 | 0.047 | 0.090 |
| InferiorTemporalGyrustemporooccipitalright | 0.078 | 0.011 | 0.000 | 0.056 | 0.100 |
| PostcentralGyrusleft | 0.126 | 0.011 | 0.000 | 0.105 | 0.148 |
| PostcentralGyrusright | 0.134 | 0.011 | 0.000 | 0.113 | 0.155 |
| SuperiorParietalLobuleleft | 0.066 | 0.011 | 0.000 | 0.045 | 0.087 |
| SuperiorParietalLobuleright | 0.076 | 0.011 | 0.000 | 0.056 | 0.097 |
| SupramarginalGyrusanteriorleft | 0.072 | 0.011 | 0.000 | 0.052 | 0.093 |
| SupramarginalGyrusanteriorright | 0.096 | 0.010 | 0.000 | 0.075 | 0.116 |
| SupramarginalGyrusposteriorleft | 0.087 | 0.011 | 0.000 | 0.066 | 0.108 |
| SupramarginalGyrusposteriorright | 0.101 | 0.011 | 0.000 | 0.080 | 0.122 |
| AngularGyrusleft | 0.081 | 0.011 | 0.000 | 0.060 | 0.101 |
| AngularGyrusright | 0.101 | 0.011 | 0.000 | 0.080 | 0.122 |
| LateralOccipitalCortexsuperiorleft | 0.135 | 0.011 | 0.000 | 0.114 | 0.156 |
| LateralOccipitalCortexsuperiorright | 0.113 | 0.011 | 0.000 | 0.092 | 0.135 |
| LateralOccipitalCortexinferiorleft | 0.124 | 0.011 | 0.000 | 0.102 | 0.145 |
| LateralOccipitalCortexinferiorright | 0.109 | 0.011 | 0.000 | 0.087 | 0.131 |
| IntracalcarineCortexleft | 0.078 | 0.011 | 0.000 | 0.057 | 0.099 |
| IntracalcarineCortexright | 0.092 | 0.011 | 0.000 | 0.071 | 0.113 |
| FrontalMedialCortexleft | 0.110 | 0.011 | 0.000 | 0.089 | 0.131 |
| FrontalMedialCortexright | 0.107 | 0.011 | 0.000 | 0.086 | 0.129 |
| JuxtapositionalLobuleCortex | 0.089 | 0.011 | 0.000 | 0.068 | 0.110 |
| JuxtapositionalLobuleCortex | 0.094 | 0.011 | 0.000 | 0.074 | 0.115 |
| SubcallosalCortexleft | 0.168 | 0.011 | 0.000 | 0.146 | 0.190 |
| SubcallosalCortexright | 0.161 | 0.011 | 0.000 | 0.139 | 0.183 |
| ParacingulateGyrusleft | 0.131 | 0.011 | 0.000 | 0.109 | 0.153 |
| ParacingulateGyrusright | 0.123 | 0.011 | 0.000 | 0.101 | 0.145 |
| CingulateGyrusanteriorleft | 0.043 | 0.011 | 0.000 | 0.022 | 0.064 |
| CingulateGyrusanteriorright | 0.048 | 0.011 | 0.000 | 0.027 | 0.069 |
| CingulateGyrusposteriorleft | 0.146 | 0.012 | 0.000 | 0.123 | 0.168 |
| CingulateGyrusposteriorright | 0.135 | 0.011 | 0.000 | 0.113 | 0.158 |
| PrecuneousCortexleft | 0.134 | 0.011 | 0.000 | 0.112 | 0.156 |
| PrecuneousCortexright | 0.137 | 0.011 | 0.000 | 0.115 | 0.159 |
| CunealCortexleft | 0.066 | 0.011 | 0.000 | 0.045 | 0.087 |
| CunealCortexright | 0.087 | 0.011 | 0.000 | 0.066 | 0.109 |
| FrontalOrbitalCortexleft | 0.178 | 0.011 | 0.000 | 0.155 | 0.200 |
| FrontalOrbitalCortexright | 0.177 | 0.011 | 0.000 | 0.155 | 0.199 |
| ParahippocampalGyrusanteriorleft | 0.152 | 0.011 | 0.000 | 0.130 | 0.174 |
| ParahippocampalGyrusanteriorright | 0.148 | 0.011 | 0.000 | 0.126 | 0.171 |
| ParahippocampalGyrusposteriorleft | 0.103 | 0.011 | 0.000 | 0.082 | 0.125 |
| ParahippocampalGyrusposteriorright | 0.086 | 0.011 | 0.000 | 0.065 | 0.107 |
| LingualGyrusleft | 0.133 | 0.012 | 0.000 | 0.110 | 0.156 |
| LingualGyrusright | 0.150 | 0.012 | 0.000 | 0.127 | 0.173 |
| TemporalFusiformCortexanteriorleft | 0.150 | 0.011 | 0.000 | 0.128 | 0.171 |
| TemporalFusiformCortexanteriorright | 0.152 | 0.011 | 0.000 | 0.130 | 0.174 |
| TemporalFusiformCortexposteriorleft | 0.154 | 0.011 | 0.000 | 0.132 | 0.176 |
| TemporalFusiformCortexposteriorright | 0.149 | 0.011 | 0.000 | 0.127 | 0.171 |
| TemporalOccipitalFusiformCortexleft | 0.087 | 0.011 | 0.000 | 0.065 | 0.108 |
| TemporalOccipitalFusiformCortexright | 0.111 | 0.011 | 0.000 | 0.090 | 0.133 |
| OccipitalFusiformGyrusleft | 0.100 | 0.011 | 0.000 | 0.078 | 0.122 |
| OccipitalFusiformGyrusright | 0.107 | 0.011 | 0.000 | 0.085 | 0.128 |
| FrontalOperculumCortexleft | 0.125 | 0.011 | 0.000 | 0.104 | 0.147 |
| FrontalOperculumCortexright | 0.098 | 0.011 | 0.000 | 0.076 | 0.119 |
| CentralOpercularCortexleft | 0.168 | 0.011 | 0.000 | 0.146 | 0.190 |
| CentralOpercularCortexright | 0.148 | 0.011 | 0.000 | 0.126 | 0.170 |
| ParietalOperculumCortexleft | 0.110 | 0.011 | 0.000 | 0.088 | 0.131 |
| ParietalOperculumCortexright | 0.107 | 0.011 | 0.000 | 0.085 | 0.129 |
| PlanumPolareleft | 0.122 | 0.011 | 0.000 | 0.100 | 0.144 |
| PlanumPolareright | 0.142 | 0.011 | 0.000 | 0.121 | 0.164 |
| HeschlsGyrusincludesH1andH2left | 0.119 | 0.011 | 0.000 | 0.097 | 0.141 |
| HeschlsGyrusincludesH1andH2right | 0.138 | 0.011 | 0.000 | 0.116 | 0.160 |
| PlanumTemporaleleft | 0.121 | 0.011 | 0.000 | 0.099 | 0.143 |
| PlanumTemporaleright | 0.130 | 0.011 | 0.000 | 0.108 | 0.152 |
| SupracalcarineCortexleft | 0.074 | 0.011 | 0.000 | 0.053 | 0.095 |
| SupracalcarineCortexright | 0.085 | 0.011 | 0.000 | 0.064 | 0.106 |
| OccipitalPoleleft | 0.091 | 0.011 | 0.000 | 0.069 | 0.113 |
| OccipitalPoleright | 0.071 | 0.011 | 0.000 | 0.049 | 0.094 |

*Note.* Standardised estimates, standard errors (SE) are reported. All model fits were $\chi^2 (17) \leq 299.229$ $p < 0.001$ , CFI $\geq 0.985$ , TLI $\geq 0.970$ , RMSEA $\leq 0.024$ , SRMR $\leq 0.015$ .

**Table S9.** Associations between *g* and subcortical volumes.

|  | Std. Est. | SE | *p* | ci.lower | ci.upper |
| --- | --- | --- | --- | --- | --- |
| lAccumbens | 0.118 | 0.011 | 0.000 | 0.096 | 0.140 |
| rAccumbens | 0.113 | 0.011 | 0.000 | 0.090 | 0.135 |
| lAmygdala | 0.084 | 0.011 | 0.000 | 0.063 | 0.105 |
| rAmygdala | 0.071 | 0.011 | 0.000 | 0.050 | 0.092 |
| lCaudate | 0.132 | 0.011 | 0.000 | 0.110 | 0.153 |
| rCaudate | 0.127 | 0.011 | 0.000 | 0.106 | 0.148 |
| lHippocampus | 0.133 | 0.011 | 0.000 | 0.111 | 0.154 |
| rHippocampus | 0.153 | 0.011 | 0.000 | 0.131 | 0.174 |
| lPallidum | 0.142 | 0.011 | 0.000 | 0.120 | 0.163 |
| rPallidum | 0.144 | 0.011 | 0.000 | 0.122 | 0.165 |
| lPutamen | 0.169 | 0.012 | 0.000 | 0.146 | 0.193 |
| rPutamen | 0.166 | 0.012 | 0.000 | 0.143 | 0.189 |
| lThalamus | 0.251 | 0.011 | 0.000 | 0.229 | 0.274 |
| rThalamus | 0.255 | 0.012 | 0.000 | 0.232 | 0.277 |

*Note.* Standardised estimates (Std. Est.), standard errors (SE) are reported with 95% confidence intervals (ci). All model fits were $\chi^2(16) \leq 209.063$ $p < 0.001$ , CFI $\geq 0.990$ , TLI $\geq 0.979$ , RMSEA $\leq 0.020$ , SRMR $\leq 0.014$ .

**Figure S1.**
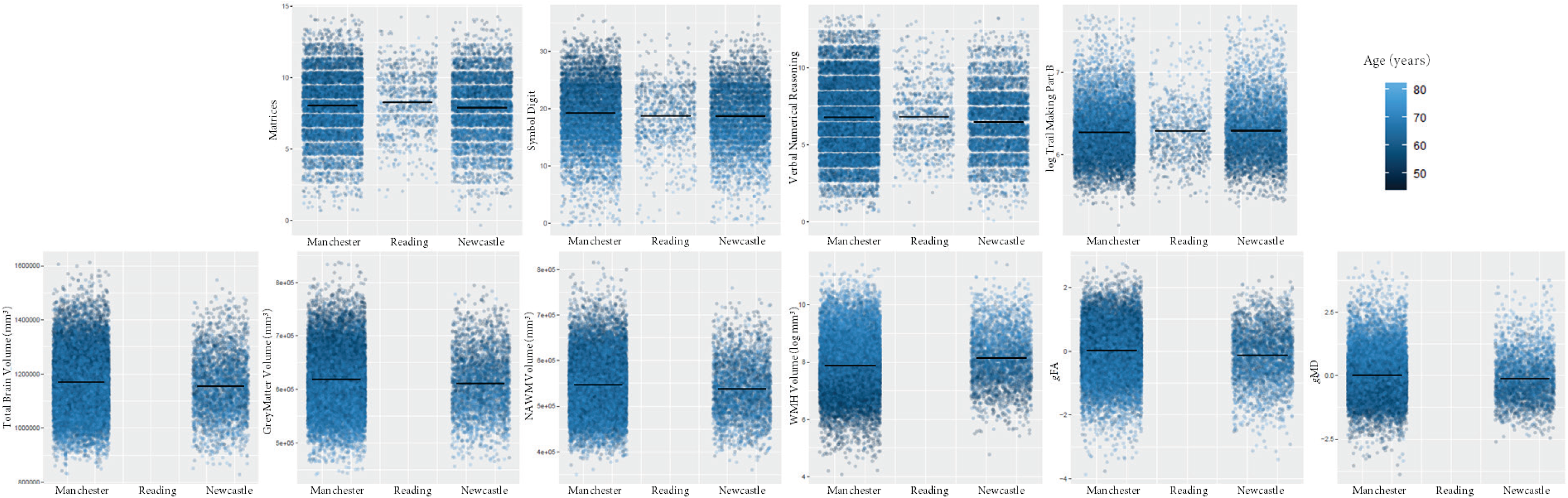
Cognitive and MRI measures across UK Biobank assessment centres. Black horizontal lines denote group means; horizontal jitter added to aid visualisation.

**Figure S2.**
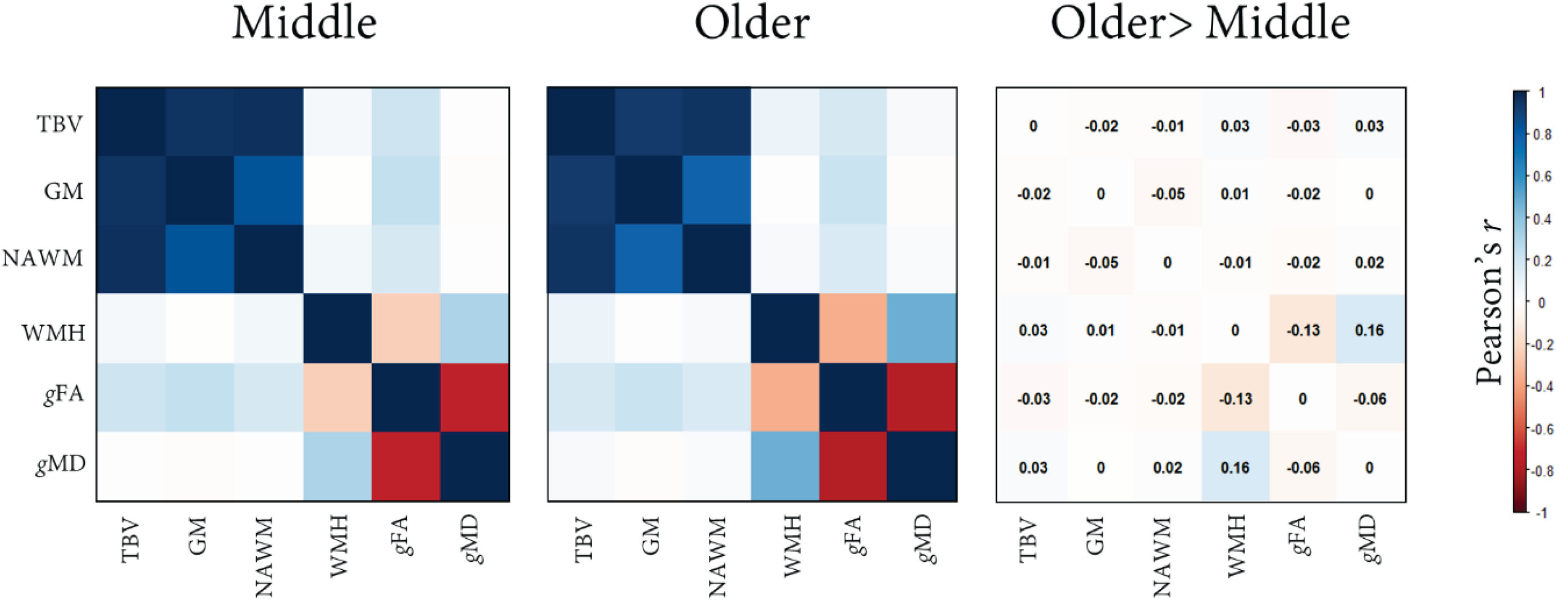
Associations among global brain images markers in middle and older age, and the difference in the magnitude of associations. TBV: total brain volume, GM: grey matter volume, NAWM: normal-appearing white matter, WMH: white matter hyperintensity volume, *g*FA: general fractional anisotropy, *g*MD: general mean diffusivity.

## Footnotes

1 We note that the splitting of the commissural fibres in this way was a pre-registered hypothesis.

